# In vivo reprogramming of murine host immune response genes following Leishmania major infection

**DOI:** 10.1101/2021.10.05.463063

**Authors:** Gopinath Venugopal, Jordan T. Bird, Charity L. Washam, Hayden Roys, Anne Bowlin, Stephanie D. Byrum, Tiffany Weinkopff

## Abstract

*Leishmania* parasites cause cutaneous leishmaniasis (CL), a pathologic disease characterized by disfiguring, ulcerative skin lesions. Both parasite and host gene expression following infection with various *Leishmania* species has been investigated in vitro, but global transcriptional analysis following *L. major* infection in vivo is lacking. Thus, we conducted a comprehensive transcriptomic profiling study combining bulk RNA sequencing (RNA-Seq) and single-cell RNA sequencing (scRNA-Seq) to identify global changes in gene expression in vivo following *L. major* infection. Bulk RNA-Seq analysis revealed that host immune response pathways like the antigen processing and presentation pathway were significantly enriched amongst differentially expressed genes (DEGs) upon infection, while ribosomal pathways were significantly downregulated in infected mice compared to naive controls. scRNA-Seq analyses revealed cellular heterogeneity including distinct resident and recruited cell types in the skin following murine *L. major* infection. Within the individual immune cell types, several DEGs indicative of many interferon induced GTPases and antigen presentation molecules were significantly enhanced in the infected ears including macrophages (*Gbp2, H2-K1, H2-Aa, H2-Ab1*), resident macrophages (*H2-K1, H2-D1, Gbp4, Gbp8, Gbp2*), and inflammatory monocytes (*Gbp2, Gbp5, Gbp7, Gbp3*). Ingenuity Pathway Analysis of scRNA-Seq data indicated the antigen presentation pathway was increased with infection, while EIF2 signaling is the top downregulated pathway followed by eIF4/p70S6k and mTOR signaling in multiple cell types including macrophages, BECs, and LECs. Altogether, this transcriptomic profile highlights known recruitment of myeloid cells to lesions and recognizes a previously undefined role for EIF2 signaling in murine *L. major* infection in vivo.

**Author summary:** *Leishmania major* cause cutaneous leishmaniasis, which is characterized by disfiguring, ulcerative skin lesions. Here, we show murine *L. major*-directed reprogramming of the host transcriptome in vivo. Our bulk RNA-Seq analyses revealed upregulation of antigen processing and presentation pathway, while the host ribosomal pathway was downregulated following *L. major* infection. Similarly, scRNA-Seq analyses revealed the upregulation of transcripts responsible for antigen presentation and host defense proteins like guanylate binding proteins (GBPs) alongside the downregulation of EIF2 signalling at the site of *L. major* infection. Overall, our transcriptomic dataset not only provides the comprehensive list of gene expression at the single-cell resolution, and highlights a previously undefined role for EIF2 signalling during murine *L. major* infection in vivo.

## Introduction

Leishmaniasis is a multifaceted disease caused by different species of obligate intracellular protozoan parasites of the genus *Leishmania*, belonging to the Trypanosomatid family. Depending on the complex interaction between the species and the host immune system, the disease can vary in severity resulting in a wide spectrum of clinical outcomes that have been classified into the following categories: cutaneous leishmaniasis (CL), mucocutaneous leishmaniasis (MCL), or diffuse CL (DCL) where symptoms remain localized to skin or mucosal surfaces, and a life-threatening condition called visceral leishmaniasis (VL) where parasites migrate to the internal organs like the liver, spleen and bone marrow(1). As per the World Health Organization (WHO), leishmaniasis is still considered to be major public health problem due to its annual incidence up to 1.7 million new cases worldwide(1, 2). *Leishmania* spp. possess a variety of virulence mechanisms, which helps parasites to survive and replicate inside the parasitophorous vacuoles of macrophages, and elimination of parasites by macrophages is critical for host resistance(3). Although, macrophages are the primary host cell for *Leishmania* parasites, recruited neutrophils(4–6) and dendritic cells(7–9) at the site of infection can also harbor parasites, suggesting that these immune cells could play an important role in host-parasite interplay. *Leishmania* parasites use several strategies to evade the host immune response for its intracellular survival including modulating the host immune response by altering T cell responses, impeding antigen display by MHCII, hindering nitric oxide production(10, 11). Importantly, *Leishmania* parasites can escape from oxidative burst and they fail to activate optimal macrophage innate immune responses(12, 13). However, changes at the transcriptional level following *Leishmania* infection within different cell types and especially within the hostile tissue microenvironment is still largely unknown.

Of the species belonging to subgenus *Leishmania*, *L. major* is an important etiological agent of cutaneous leishmaniasis (CL) and possesses clinical and epidemiological importance, especially in parts of Asia, the Middle East, Northern Africa, and Southern Europe(14). Although CL is not fatal and considered to be a self-healing disease, the development of nodules or papules followed by ulcerations at the site of promastigote infection is the hallmark of the disease; importantly, both parasite replication and the host immune response can contribute to the disease pathology(14). CL produces permanently disfiguring skin lesions that are associated with massive immune cell recruitment, across the blood vascular endothelium, and into the skin where the parasite resides(15–19). While CD4^+^ Th1 cells producing IFNγ are required to activate macrophages to kill parasites, the exacerbated activation and sustained recruitment of immune cells including neutrophils, NK cells, Ly6C^+^ inflammatory monocytes, and CD4^+^ and CD8^+^ T lymphocytes induces a chronic inflammatory response; this chronic inflammation leads to tissue necrosis and skin damage, a feature of non-healing lesions(20–22).

The *L. major* genome was completed in 2005(23). Until recently, many studies examining the host response relied mostly on microarray-based or serial analysis of gene expression tag approaches. Using these approaches, previous studies compared the host gene expression profile between infection with promastigotes and amastigotes or different species of *Leishmania*(24–30). Transcriptomic studies dramatically enhanced our understanding of CL. In general, these studies concluded that human or murine macrophages infected by various *Leishmania* species downregulate pro-inflammatory gene expression, while concomitantly upregulating the expression of anti-inflammatory genes(31, 32). However, microarray-based approaches have technical limitations such as hybridization and cross hybridization artefacts, dye-based detection issues, certain probes cannot be included on the microarrays, and the inability to detect 5’ and 3’ UTRs boundaries(33, 34). In recent years, RNA-Seq has emerged as a powerful tool to study transcriptional changes in many disease conditions due to its high sensitivity. Moreover, transcriptomic profiling using RNA-Seq following infection with various species of *Leishmania* has been mostly applied to in vitro experiments and some studies have investigated transcriptional changes in the human or murine host as well as the parasite simultaneously(35–42). A recent study comparing the gene expression profile of primary cutaneous lesions from *L. braziliensis*-infected patients with or without pentavalent antimony treatment revealed most of the differentially expressed transcripts were correlated with components of cytotoxicity related pathways and parasite load in the skin(39). While another study using RNA-Seq from the same group revealed a consistent and significant myeloid interferon stimulated gene (ISG) signature in skin lesions from *L. braziliensis*-infected patients(43). Altogether, these studies revealed that the host immune response upregulates transcripts related to both pro-inflammatory and anti-inflammatory responses during leishmaniasis(44, 45). Furthermore, none of the RNA-Seq studies to date have characterized the global transcriptional reprogramming following *L. major* infection in vivo, which is the most widely used murine model of CL to study disease pathogenesis and parasite-host interactions. The *Leishmania* field also lacks a comprehensive murine transcriptomic profile that applies more recent genomic technologies like scRNA-Seq to identify transcriptomic changes within individual cell types in vivo. Therefore, the purpose of this study was to generate comprehensive transcriptomic profile using a combination of bulk RNA-Seq and scRNA-Seq to identify global changes in gene expression that occur during murine *L. major* infection in vivo.

Our bulk RNA-Seq transcriptomic analyses revealed the antigen presentation pathway was significantly upregulated while ribosomal pathways were significantly downregulated by the host following *L. major* infection. Our scRNA-Seq analyses revealed a cellular heterogeneity including distinct resident and recruited cell types at the site of *L. major* infection. Confirming the bulk RNA-Seq, we found macrophages, blood endothelial cells (BECs), and lymphatic endothelial cells (LECs) display a transcriptomic profile associated with increased antigen presentation and decreased EIF2, eIF4/p70S6K, and mTOR signalling suggesting cells in lesions are undergoing a stress response while they participate in Th1 protective immunity. Overall, this study combines bulk RNA-Seq and scRNA-Seq to assemble a comprehensive dataset that defines how the murine host response reprograms individual cell types following *L. major* infection to combat the infection.

## Results

### Mouse model of *L. major* infection

To characterize the transcriptomic landscape during *L. major* infection in vivo, we employed next-generation genome sequencing on RNA analysis with ears from mice that were infected with *L. major* promastigotes and uninfected naive control ears. During this experimental murine model of *L. major* infection, lesion volume peaks between 4-6 weeks post-infection (p.i.) and then the lesion resolves spontaneously, and parasites are present in lesions at this time point (Fig 1A-B). As a result, the host transcriptome was investigated from ear samples collected from experimentally-infected mice at 4 weeks p.i. and compared to naive controls.

**Figure 1.**
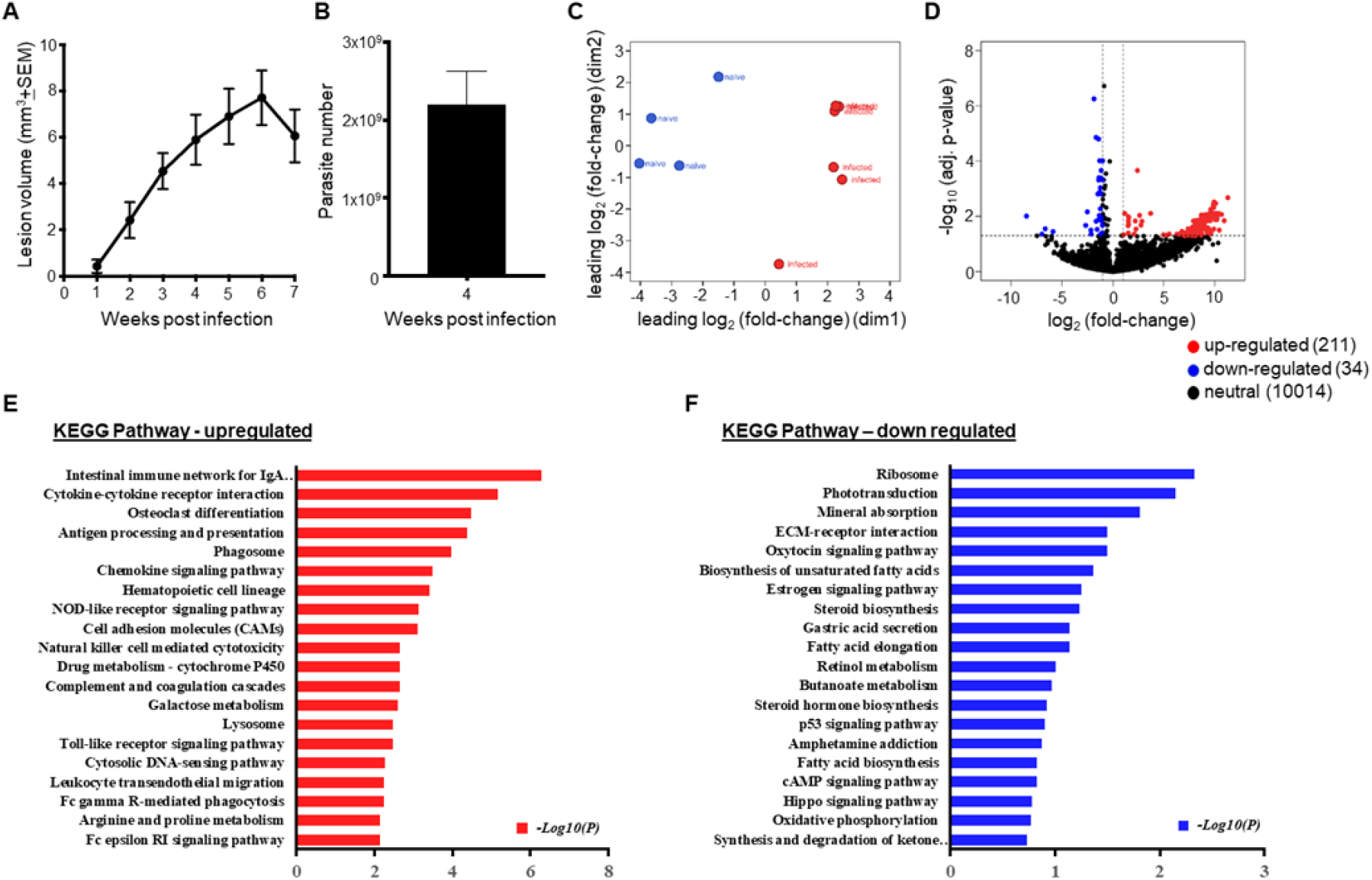
*Leishmania* infection is associated with differential regulation of host immune response pathways in vivo. **(A)** C57BL/6 mice were infected with 2x10^6^ *L. major* metacyclic promastigote parasites intradermally in the ear and lesion development was monitored over time. Data are pooled from 4 experiments (n=30) and shown as +SEM. **(B)** At 4 week post-infection, parasite numbers in the ear were quantified. Data are pooled from 2 experiments (n=20) and shown as +SEM. **(C)** Multi- Dimensional Scaling (MDS) plot showing the gene expression profile between naive and *L. major-*infected ears. **(D)** Volcano plot of all DEGs in naive and infected ears. Red dots represent upregulated DEGs with log2FC >1 and p-value <0.05. Blue represents down-regulated DEGs with log2FC <−1 and p-value <0.05. Only annotated genes are shown in plot. FC, fold-change. **(E-F)** Bulk RNA-Seq analysis indicates DEGs that are highly correlated with signaling pathways. KEGG enrichment analysis of top 20 upregulated (E), and top 20 downregulated (F) pathways enriched among the DEGs between naive and *L. major*-infected ears (pathways selected by significant and FC > 1.5, list includes rank, significance and adjusted average).

### Enriched pathways for differentially expressed genes during *L. major* infection by bulk RNA-Seq

To compare the gene expression profiles of infected and naive mice, bulk RNA-Seq was performed on *L. major*-infected ears and naive control ears. Transcriptional analysis revealed that ears from infected mice and naive controls were distinct from one another, as determined by multidimensional scaling (MDS) plot and DEG analysis. MDS plot shows the positions of each sample, with samples from different experimental groups being well separated, and samples from the same experimental group clustering together (Fig 1C). Therefore, the distance between samples reveals the distinct pattern of gene expression between the infected and naive animals (Fig 1C). To investigate transcriptomic signatures associated with infection, we carried out DEG analysis between infected mice and naive controls by comparing the RNA-Seq read counts of the various genes and subsequently applying the cut-off criteria. High and low expression genes (logCPM><1) were included in the volcano plot showing transcriptional differences observed between infected and naive ears (Fig 1D). The gene expression profiles derived from the RNA-Seq data were calculated using the RPKM method and a fold change >2 and p<0.05 were considered statistically significant. Of more than 10,800 genes that were detectable in the infected ears, we observed that 211 genes were upregulated and 34 genes were downregulated, while 10,014 genes did not show any significant differences between naive and *L. major*- infected ears (Fig 1D).

Gene set enrichment analysis (GSEA) using KEGG pathways revealed a total of 276 enriched pathways which includes pathways involved in both disease conditions and molecular signaling networks. Specifically, the antigen processing and presentation pathway was found to be significantly enriched amongst DEGs, while the ribosomal pathway was significantly downregulated during *L. major* infection (Fig 1E- F). In addition to antigen processing and presentation, we observed many other host immune response pathways upregulated with infection including: cytokine-cytokine receptor interaction, phagosome, chemokine signaling, cell-adhesion molecules pathway, NK cell mediated cytotoxicity, leukocyte trans-endothelial migration and Fcγ receptor-mediated phagocytosis (Fig 1E). Conversely, top 20 downregulated pathways enriched for DEGs in *L. major* infection were related to ribosomal translation, mineral absorption, ECM-receptor interaction, biosynthesis of unsaturated fatty acids, and steroid biosynthesis (Fig 1F). The top 10 KEGG pathways for both upregulated and downregulated pathways with the fold change and adjusted ‘*p*’ value are listed in Table 1. A number of disease-specific KEGG pathways appeared prominent in the enrichment analysis including *Staphylococcus aureus* infection, autoimmune thyroid disease, and graft vs. host disease (Table S1). Importantly, leishmaniasis emerged as one of the top disease pathways highlighting the quality of the input data for the analysis (Table S1).

**Table 1.** List of top 10 KEGG pathways enriched for differentially expressed genes (DEGs) following L. major infection.

| Pathway regulation | KEGG enriched pathways | Avg. log-fold change | Adj. p value |
| --- | --- | --- | --- |
| Up regulated | Intestinal immune network for IgA production | 5.23228 | 5.08E-07 |
|  | Cytokine-cytokine receptor interaction | 8.337469 | 6.97E-06 |
|  | Osteoclast differentiation | 7.623207 | 3.31E-05 |
|  | Antigen processing and presentation | 4.468822 | 4.21E-05 |
|  | Phagosome | 6.527853 | 0.000106 |
|  | Chemokine signaling pathway | 8.329182 | 0.000328 |
|  | Hematopoietic cell lineage | 6.848572 | 0.000381 |
|  | NOD-like receptor signaling pathway | 6.638004 | 0.000744 |
|  | Cell adhesion molecules | 6.486765 | 0.000769 |
|  | Natural killer cell mediated cytotoxicity | 8.022157 | 0.002282 |
| Down-regulated | Ribosome | -0.696061 | 0.004705 |
|  | Phototransduction | -0.907101 | 0.007031 |
|  | Mineral absorption | -0.992836 | 0.015538 |
|  | ECM-receptor interaction | -0.889139 | 0.031673 |
|  | Oxytocin signaling pathway | -0.948893 | 0.031673 |
|  | Biosynthesis of unsaturated fatty acids | -1.474018 | 0.042932 |
|  | Estrogen signaling pathway | -1.199847 | 0.055815 |
|  | Steroid biosynthesis | -0.908215 | 0.058351 |
|  | Gastric acid secretion | -0.907101 | 0.072276 |
|  | Fatty acid elongation | -0.892548 | 0.072946 |

### Differential gene expression in immune-related pathways during *L. major* infection by bulk RNA-Seq

Using hierarchical clustering analysis, we found that a large number of genes were robustly induced in the infected ears compared to the naives (Fig 2A-D). A heat map of the DEGs shows the expression profiles of infected and naive mice resulted in separate clusters (Fig 2). Hierarchical clustering reveals the host immune response to *L. major* infection is closely linked with DEGs from the antigen processing and presentation pathway (*Cd4, H2-Q6, H2-M3, H2-Q4, Ifng, Cd8b1, H2-T22, Rfx5, Tap1, H2-Q7*), chemokine signaling (*Cxcl9, Ccl5, Ccr7, Cxcl10, Cxcl5, Cxcl16, Cxcl1, Fgr, Pik3cd*), and cell adhesion molecules (*Cd4, Itgam, Itgal, Ctla4, Icos Itga6, Cd274, Cd28, Cd86, Selplg, Vcam1*) (Fig 2A-C). Additionally, other immune network pathways enriched for DEGs included cytokine-cytokine receptor interaction, phagosome, toll-like receptor signaling, and leukocyte trans-endothelial migration pathway (Fig S1A-D). In contrast, the biological processes downregulated with infection include ribosomal biogenesis (*Rpl3, Rpl37, Rps5, Rpl11, Rplp1, Rpl28, Rpl19, Rps28, Rps14*) (Fig 2D). Of note, infected mice clustered together for antigen processing and presentation, chemokine signaling, and the cell adhesion molecules pathways (Fig 2A- C), but one mouse in each group clustered with the opposing experimental group for the ribosomal pathway (Fig 2D). Overall, these results demonstrate the host transcriptome undergoes reprogramming in the skin during *L. major* infection.

**Figure 2:**
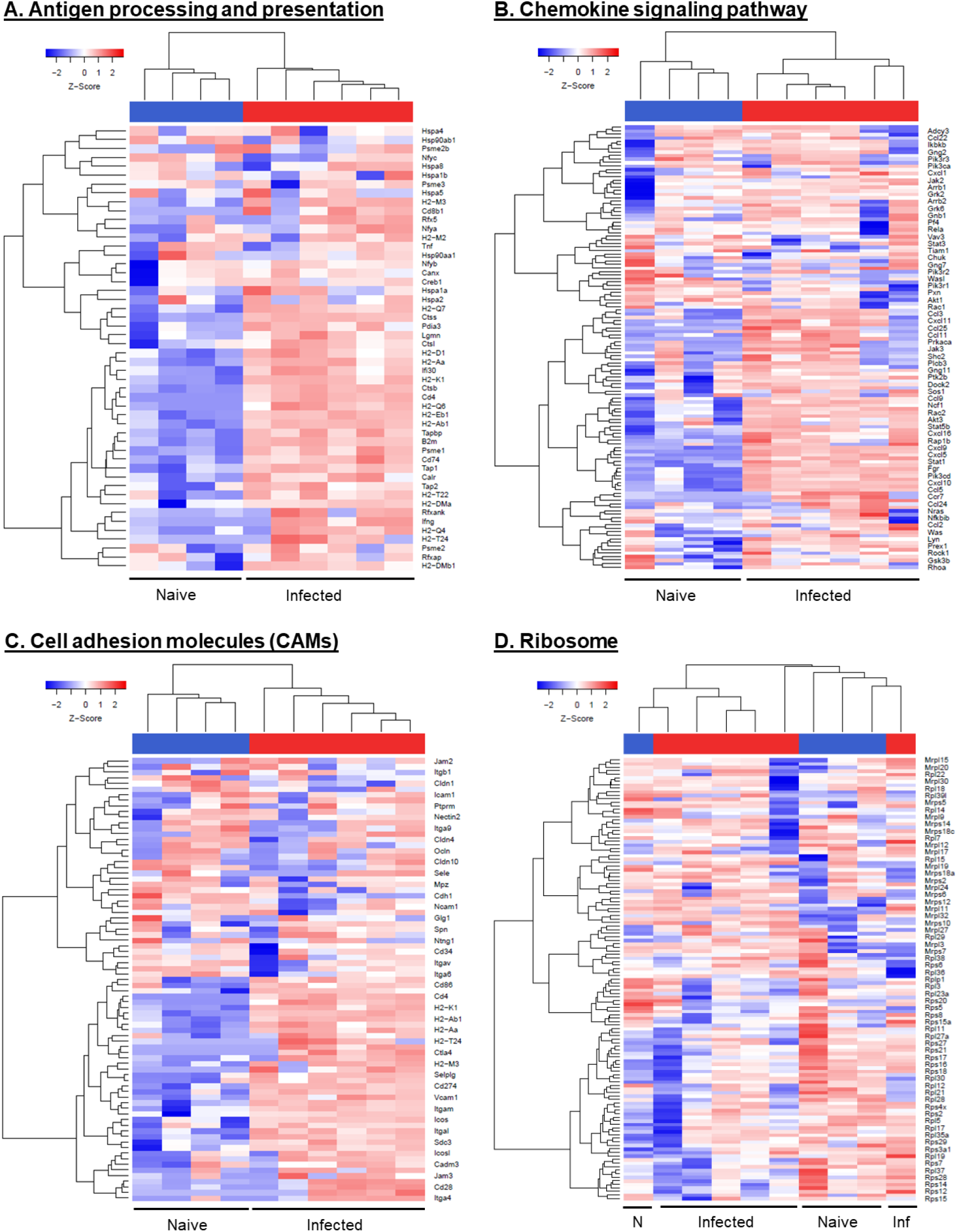
Heat map analysis of host transcriptional responses to *L. major* infection in vivo. Heat maps of DEGs in infected ears compared to naive controls. The DEGs involved in the host immune response pathways by KEGG enrichment analysis whether upregulated **(A, B, and C)** or downregulated **(D)** in the infected ears presented as heat maps. Hierarchical clustering of the expression profile was grouped according to functional categories. Heat maps indicate the FC of gene expression in *L. major*-infected ears >2-fold (red) or <2-fold (blue).

### scRNA-Seq reveals the cellular heterogeneity and altered transcriptomic profile of individual cell types during murine *L. major* infection in vivo

The bulk RNA-Seq analysis revealed global changes in the transcriptional profile between infected mice and naive controls following *L. major* inoculation. To further investigate transcriptomic changes within individual cell types present in leishmanial lesions, scRNA-Seq was performed to provide a deeper understanding of how individual cells function in the tissue microenvironment. Single cells from the ears of infected and naive mice were bar-coded and sequenced using the droplet-based 10X Genomics Chromium platform (Fig 3A). After quality control assessment and filtering, the datasets were processed using Cell Ranger software. Unbiased hierarchical clustering using Seurat provides single-cell transcriptional profiling with 26,558 cells and displayed the cellular heterogeneity which includes both resident and recruited cell types. Cell populations from 35 distinct cell types were defined using canonical markers from published literature and online databases (Fig 3B)(46). The dot plot representing the cell type-specific canonical markers for each cell lineage used to distinguish the 35 distinct clusters is provided (Fig 4A). Amongst the 35 cell types, we identified 16 cell types containing immune cells. Feature plots show the expression of cell type-specific canonical markers (in addition to the cell type-specific canonical markers in Fig 4A) for 12 clusters with corresponding cell types (Fig 4B). Additionally, a heatmap shows the canonical cell type markers from all the immune cell types along with blood endothelial cells (BECs) and lymphatic endothelial cells (LECs) (Fig S2).

**Figure 3.**
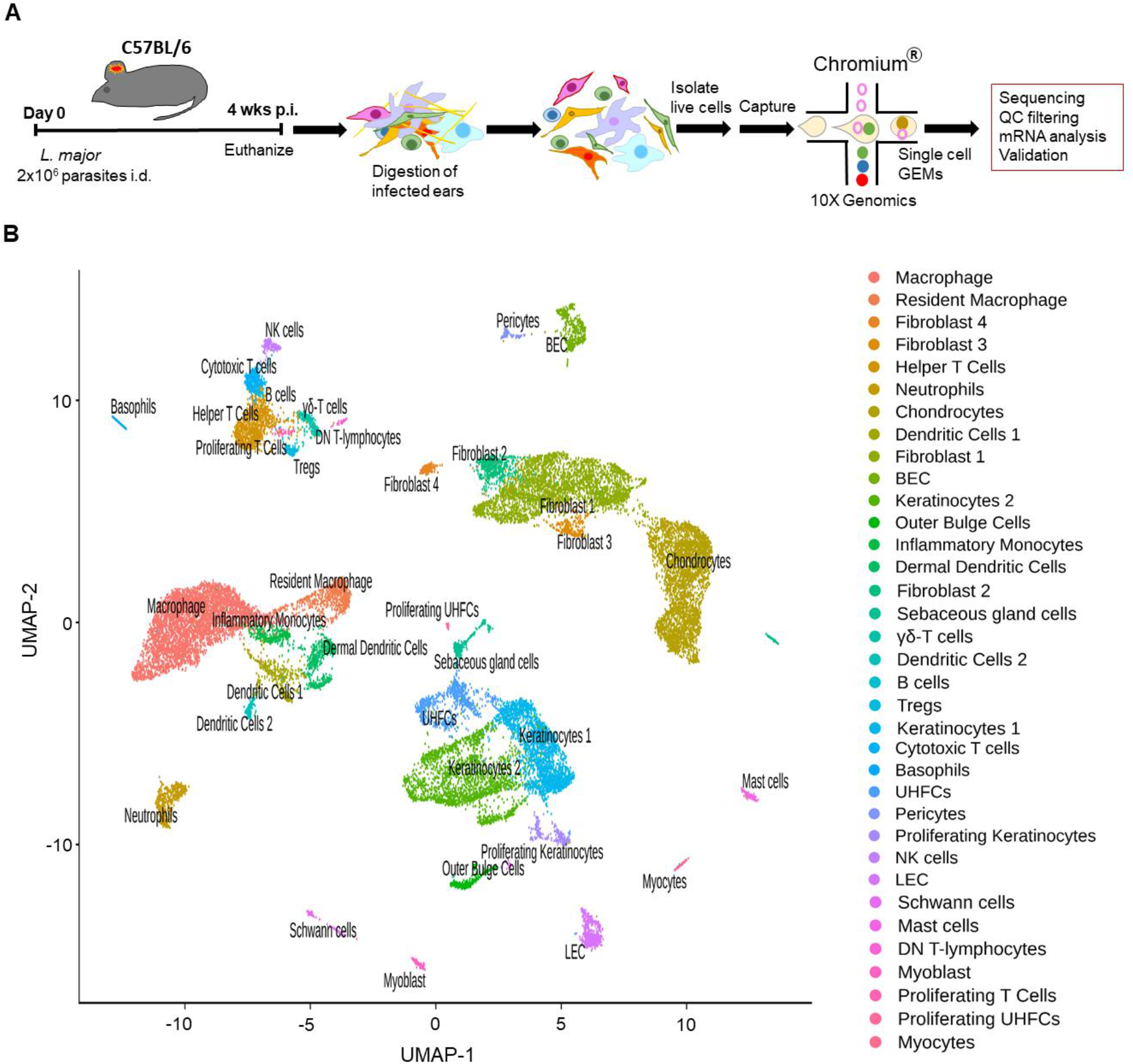
scRNA-Seq reveals cellular heterogeneity including distinct resident and recruited cell types in the skin following *L. major* infection. (A) C57BL/6 mice were infected or not with *L. major* parasites in the ear, and ears were digested to isolate RNA for scRNA-Seq analysis. Schematic of cell isolation, cell processing, capture by droplet-based device. **(B)** Uniform Manifold Approximation and Projection (UMAP) plot revealed cellular heterogeneity with 35 distinct clusters of cells identified and color-coded (both naive and infected groups combined). Seurat’s FindClusters function was used to identity each cell cluster and cell type designation is located to the right.

**Figure 4:**
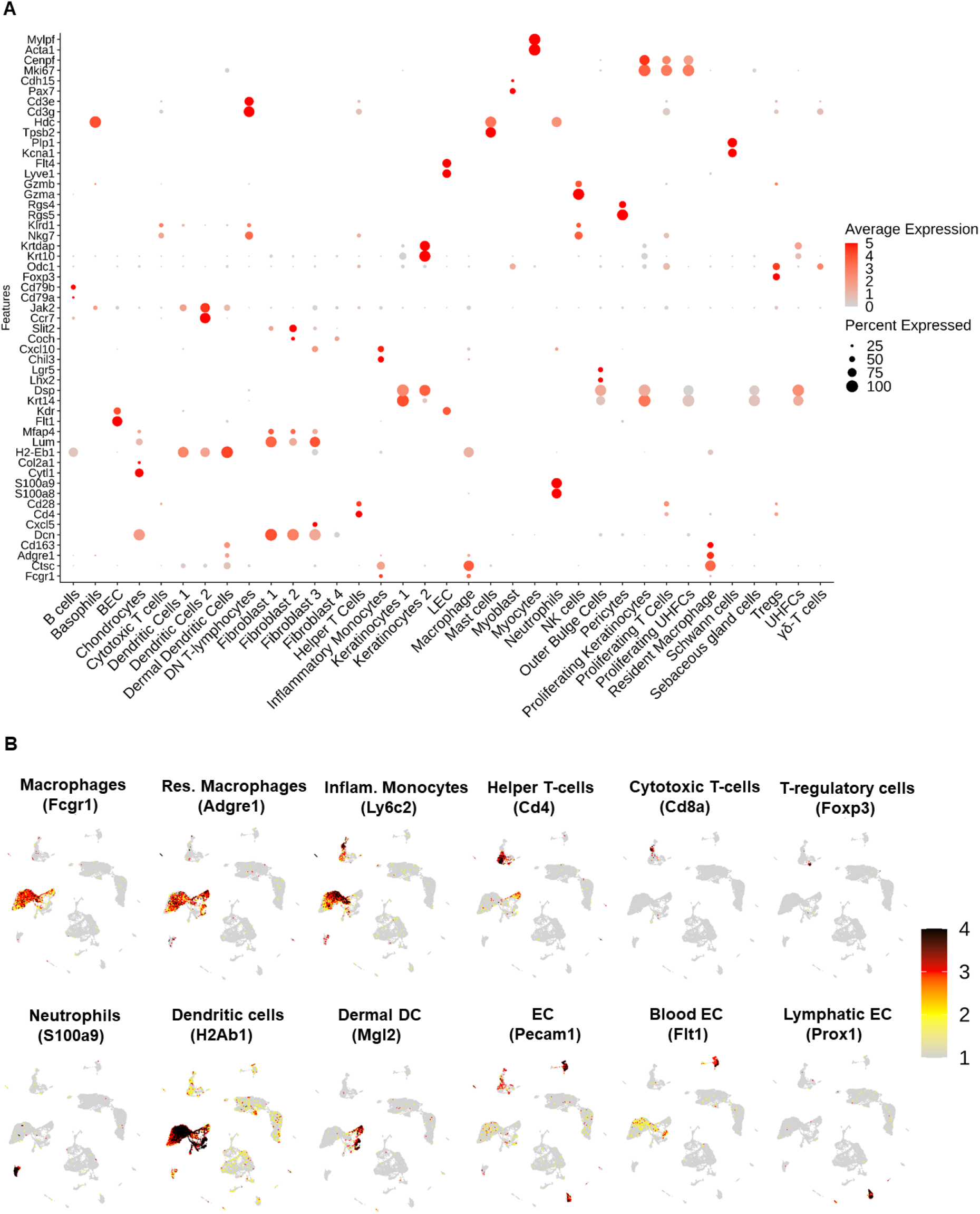
Cell type identification and cluster-specific gene expression. **(A)** Relative expression of 35 different cell type cluster-specific genes shown as dot plots with two genes per cluster. Dot size indicates the percentage of cells expressing each gene, the dot color indicates the expression level, and ordering is performed from low to high expressing cells. **(B)** Feature plots of expression distribution for selected cluster-specific genes used to define the cell types. Expression levels for each marker is color-coded and overlaid onto UMAP plot. Cells with the highest expression level are colored black.

### Detection of *Leishmania major* transcripts in multiple cell types other than macrophages

Additionally, we aligned the reads to *Leishmania major* (LM) reference genome to detect the presence of *Leishmania* transcripts in 35 different cell type clusters. Interestingly, the differential expression of LM transcripts from scRNA-Seq revealed 20 of 35 cell types have at least one LM transcript within that cell. As predicted, we found macrophages are the top immune cell type expressing LM transcripts with about 10% of cells containing LM transcripts (Fig 5A-B). We found at least 2% of cells contain LM transcripts in other immune cell types such as resident macrophages, DCs, and neutrophils. At least 1% of the cells in CD4^+^ Th cells, CD8^+^ cytotoxic T cells, T regulatory cells, and basophils also possessed detectable LM transcripts (Fig 5A-B). Consistent with previous findings, we found fibroblasts and keratinocytes also harbor LM transcripts(47). Surprisingly, we also detected LM transcripts in >5% myoblasts but not myocytes and almost 1% of the BECs. Overall, our data shows that multiple other cell types at the infection site harbors LM transcripts suggesting other cell types maybe infected with parasites alongside macrophages, which is the well-established primary host cell for *Leishmania* parasites.

**Figure 5:**
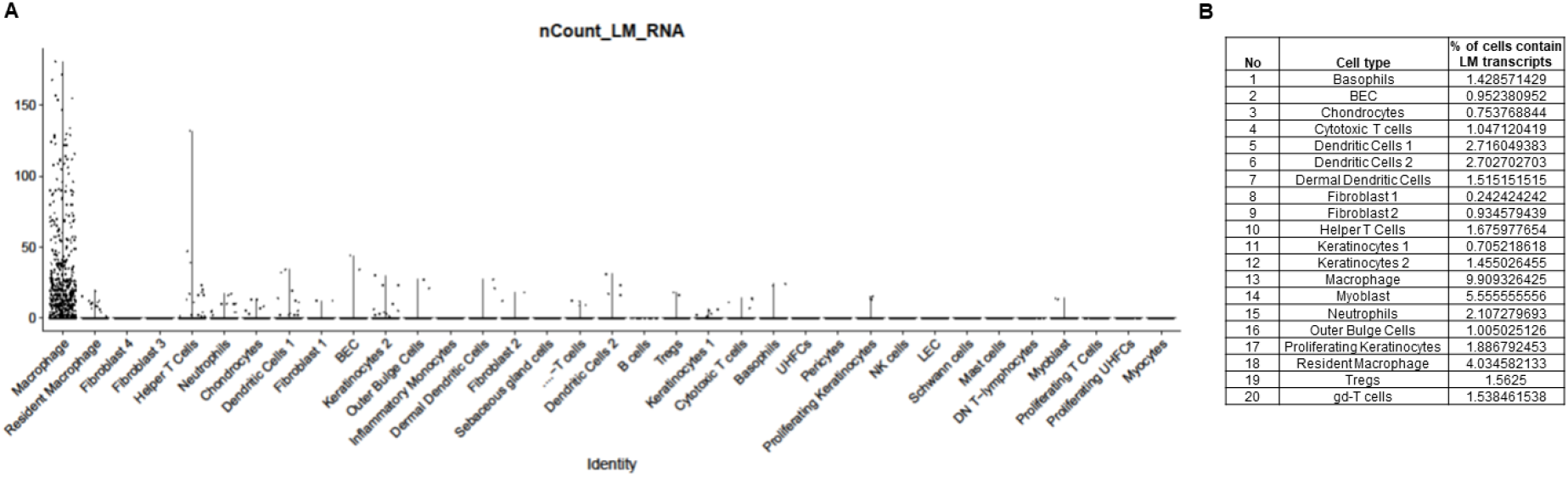
Presence of *L. major* transcripts in multiple cell types. **(A)** Differential expression of *L. major* parasite transcripts in 35 different cell types. Number of cells containing parasite transcripts was shown in violin plots. **(B)** Table summarizes the percentages of individual cell that contain *L. major* transcripts.

### scRNA-Seq confirms the immune cell recruitment at the site of *L. major* infection in vivo

Murine *L. major* infection leads to the recruitment of immune cells such as inflammatory monocytes, neutrophils, and macrophages to the site of infection. Particularly macrophages play roles in housing the parasite as the replicative niche for the pathogen, as well as a role in parasite control by killing the pathogen. BECs mediate immune cell recruitment to the infected and inflamed tissue and LECs promote immune cell migration away from the infected skin. Therefore, we speculate that immune cells and ECs participate in parasite control and/or immunopathology during CL. As a result, the remainder of the study focuses on 7 immune cell types along with the BEC and LEC clusters. Our UMAP projection displays 13,034 cells in naive ears and 13,524 cells in the infected ears (Fig 6A). Consistent with previous findings in CL, the UMAP plot confirms a significant recruitment of various immune cell types such as inflammatory monocytes, neutrophils, macrophages, dendritic cells, NK cells, and CD4^+^ and CD8^+^ T cells in the infected ears that are seen at higher frequencies compared to naive controls (Fig 6A). Concordant with our scRNA-Seq results, flow cytometric analysis detected a significant increase in the frequency and cell number of macrophages (Fig 6B-C), Ly6C^+^ inflammatory monocytes (Fig 6D-E) and neutrophils (Fig 6F-G) in infected ears compared to naive controls, while no significant alterations in the BEC or LEC populations were observed (Fig 6H-J). Altogether, these data confirm the enhanced immune cell migration during *L. major* infection and transcriptional changes within these individual cell types were investigated.

**Figure 6:**
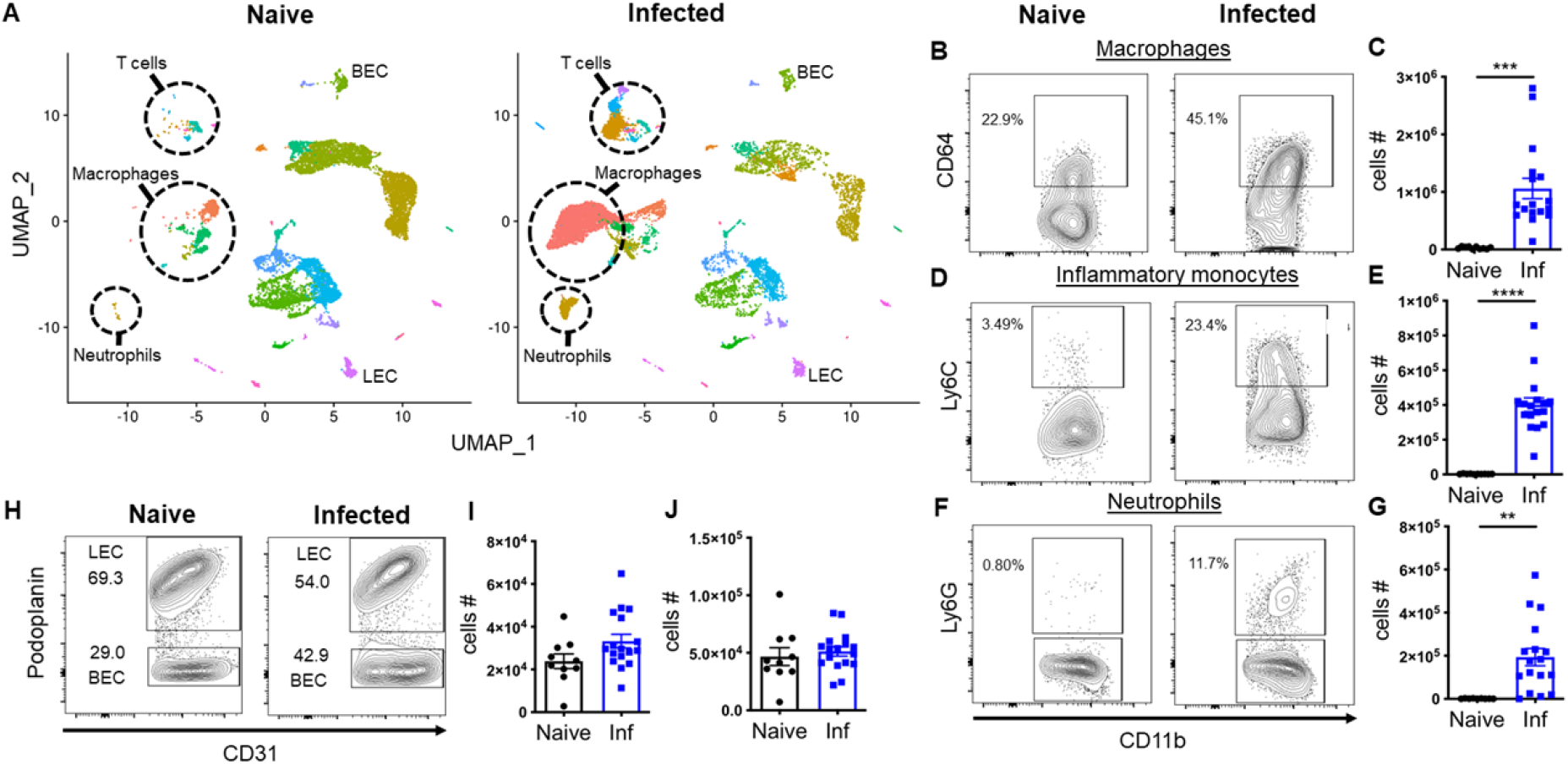
scRNA-Seq analysis reveals an enhanced immune cell recruitment to the inflamed tissue during *L. major* infection. **(A)** scRNA-Seq UMAP plots of naive and infected ears showing resident and recruited cell populations/clusters. **(B, D, F)** Representative flow cytometry plots showing the percentage of CD64^+^ macrophages (B), Ly6C^+^ inflammatory monocytes (D), and Ly6G^+^ neutrophils (F) of naive and infected ears at 4 week p.i. Cells were gated on total, live, singlets, CD45^+^ CD11b^+^ cells. **(C, E, G)** Cell numbers of CD64^+^ macrophages (C), inflammatory monocytes (E), neutrophils (G) from naive and infected ears. **(H-J)** ECs were gated on total, live, singlets, CD31^+^ cells. Dermal BECs and LECs were separated by podoplanin expression during FACS analysis. (H) Representative flow cytometry dot plots showing the percentages of BECs and LECs from naive and infected ears 4 week p.i. Corresponding cell numbers of BECs (I) and LECs (J) from naive and infected ears. Data are representative of at least two independent experiments involving 10-17 mice. Data are presented as mean ±SEM. \*\**p < 0.005*, \*\*\**p < 0.0005*, \*\*\*\**p < 0.0001*, unpaired *t*-test.

### Differential gene expression of immune cell types during *L. major* infection

To explore the transcriptional changes in a cell type-specific manner, the DEGs were compared between infected and naive mice within an individual cell type following *L. major* infection. A volcano plot showing DEGs for macrophages, resident macrophages, inflammatory monocytes, and neutrophils reveals several markers indicative of an increase in myeloid cells in leishmanial lesions (Fig 7A-D). For instance, transcripts commonly elevated within the top 10 DEGs in myeloid cells and dendritic cells (DCs) include *B2m, H2-K1, Gbp2, ligp1,* whereas multiple ribosomal proteins were significantly downregulated within the top 10 DEGs among myeloid cells and DCs (Table 2, 3, and Fig S3). We found consistent elevation of various interferon-induced GTPases like guanylate binding protein (GBP) transcripts with *L. major* infection in macrophages (*Gbp2*) (Table 2A), resident macrophages (*Gbp4, Gbp8, Gbp2*) (Table 2B), inflammatory monocytes (*Gbp2, Gbp5, Gbp7, Gbp3*) (Table 3A), and DCs (*Gbp2*) (Fig S3). Many of the transcriptomic differences detected in myeloid cells, such as elevated GBPs, were also found in EC populations. For instance, BECs expressed increased *Gbp4* and *Gbp2*, and LECs expressed increased *Gbp4, Gbp2,* and *Gbp7* upon infection (Table 4A-B). Furthermore, we detected a significant elevation of both MHCI and MHCII molecules in the infected ears in myeloid cells and ECs, which include *H2-K1, H2-Aa, H2-Ab1* in macrophages; *H2-K1, H2-D1* in resident macrophages; *H2-K1* in DCs, *H2-K1, H2-Aa, H2-Ab1* in BECs; and *H2-K1, H2-D1, H2-Q7, H2-Aa, H2-Ab1* in LECs (Table 2, 4, and Fig S3). In contrast to myeloid cells, DCs, and ECs, we only detected few transcripts that are significantly elevated with infection in T cells including *B2m, Satb1, Gm42418, Gimap6* in CD4^+^ T cells and *Gm42418* in CD8^+^ T cells (Fig S4).

**Figure 7:**
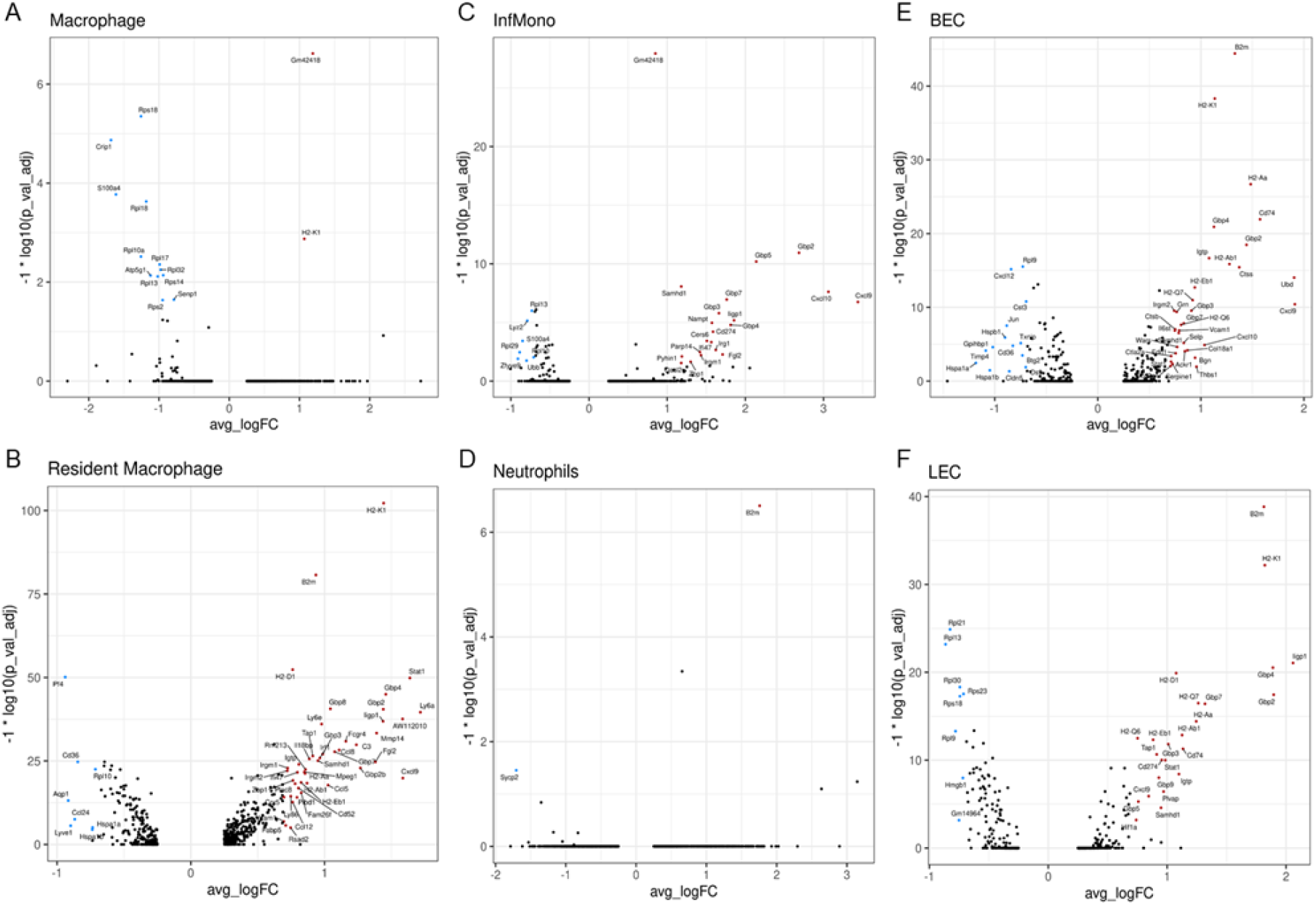
Differentially expressed genes in selected immune cell types during *L. major* infection. Volcano plot showing the DEGs in macrophages (A), resident macrophages (B), inflammatory monocyte (C), neutrophils (D), BECs (E), and LECs (F). Colored dots indicate genes at least 2 (natural log ∼0.693) fold increased (red) or decreased (blue) in infected cells relative to naive cells with an adjusted p-value < 0.05.

**Table 2.** List of top 10 DEGs enriched in macrophages and resident macrophages following L. major infection.

**Macrophages**
| Gene name | Avg. log-fold change | Adj. p value |
| --- | --- | --- |
| **<u>Upregulated</u>** |  |  |
| *H2-K1* | 1.203478 | 2.59E-30 |
| *Gm42418* | 1.013645 | 3.48E-30 |
| *B2m* | 0.798061 | 3.33E-28 |
| *AW112010* | 2.205139 | 1.15E-26 |
| *Fth1* | 1.136905 | 2.54E-26 |
| *Cxcl16* | 1.778107 | 5.99E-22 |
| *Gbp2* | 1.141379 | 1.24E-21 |
| *H2-Aa* | 1.205499 | 1.62E-20 |
| *H2-Ab1* | 1.264542 | 9.46E-20 |
| *Mmp14* | 1.784564 | 6.05E-19 |
| **<u>Down regulated</u>** |  |  |
| *Rpl32* | -1.2065 | 3.25E-34 |
| *Rpl13* | -1.23205 | 2.53E-33 |
| *Lyz2* | -1.48379 | 5.03E-32 |
| *Rpl18* | -1.22252 | 4.01E-30 |
| *S100a4* | -1.6294 | 6.25E-30 |
| *Rps18* | -1.21541 | 1.51E-28 |
| *Rps14* | -1.0183 | 1.73E-24 |
| *Fau* | -1.03269 | 6.54E-23 |
| *Ifitm3* | -1.30983 | 8.25E-23 |
| *Tpt1* | -0.83138 | 1.52E-22 |

**B** **Resident macrophages**
| Gene name | Avg. log-fold change | Adj. p value |
| --- | --- | --- |
| **<u>Upregulated</u>** |  |  |
| *H2-K1* | 1.441768 | 6.11E-103 |
| *B2m* | 0.935358 | 1.97E-81 |
| *H2-D1* | 0.761557 | 4.47E-53 |
| *Stat1* | 1.637833 | 1.41E-50 |
| *Gbp4* | 1.457725 | 1.05E-45 |
| *Gbp8* | 1.043486 | 2.40E-41 |
| *Gbp2* | 1.43904 | 3.27E-41 |
| *Ly6a* | 1.716371 | 2.56E-40 |
| *AW112010* | 1.583531 | 2.66E-38 |
| *ligp1* | 1.437197 | 1.31E-37 |
| **<u>Down regulated</u>** |  |  |
| *Pf4* | -0.93903 | 7.66E-51 |
| *Cd36* | -0.8449 | 1.88E-25 |
| *Rpl12* | -0.64508 | 1.92E-25 |
| *S100a6* | -0.59164 | 2.65E-24 |
| *Rps25* | -0.64782 | 2.88E-24 |
| *Rpl10* | -0.71332 | 3.09E-23 |
| *Rnf213* | 0.85734 | 3.50E-23 |
| *Rps12* | -0.5271 | 3.76E-23 |
| *Rpl30* | -0.60024 | 4.48E-23 |
| *Rps18* | -0.56378 | 9.59E-21 |

**Table 3.**
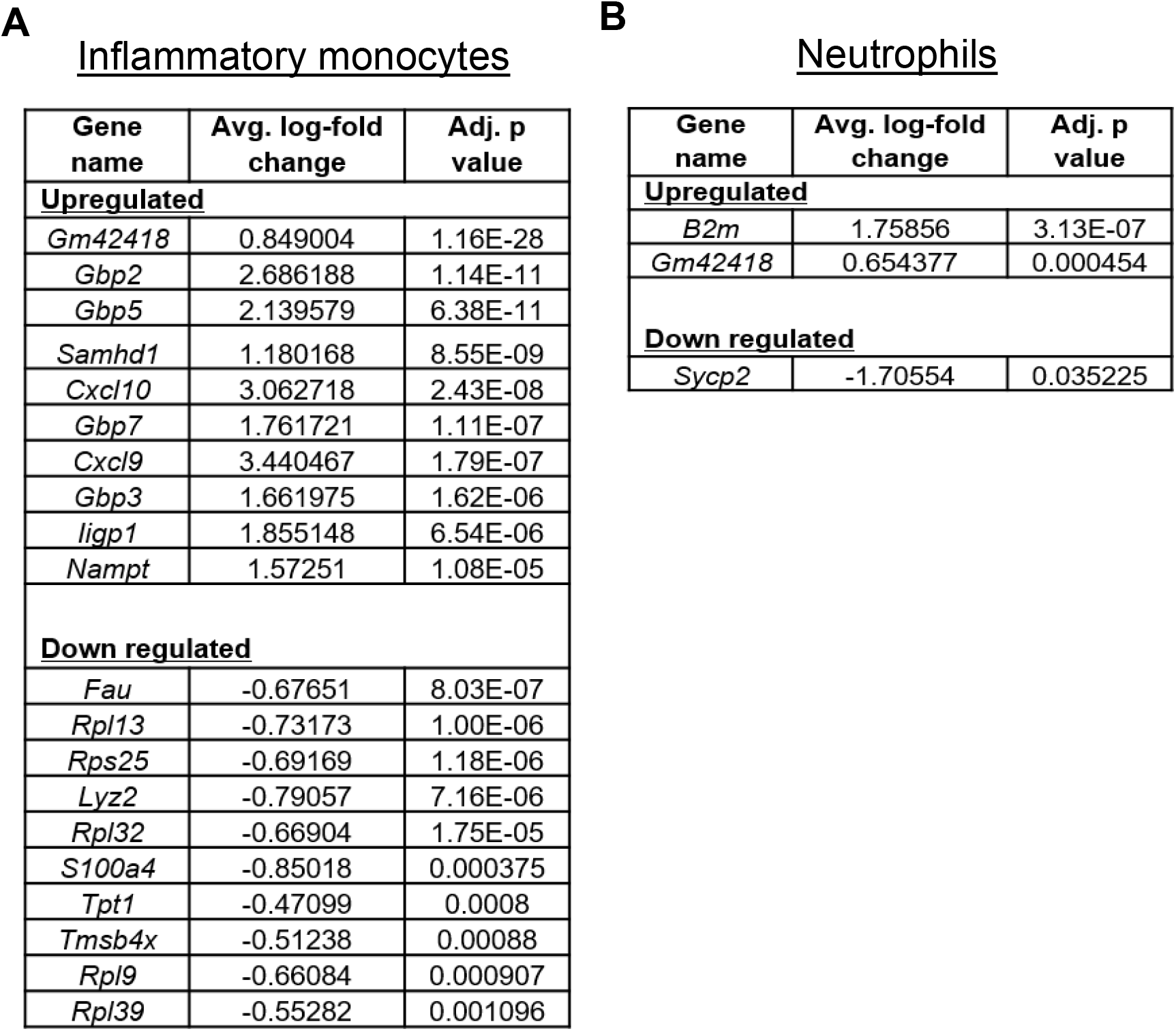
List of top 10 DEGs enriched in inflammatory monocytes and neutrophils clusters following *L. major* infection.

**Table 4.** List of top 10 DEGs enriched in BEC and LEC clusters following L. major infection.

| Gene name | Avg. log-fold change | Adj. p value |
| --- | --- | --- |
| **Upregulated** |  |  |
| *B2m* | 1.330157 | 3.83E-45 |
| *H2-K1* | 1.134934 | 4.90E-39 |
| *H2-Aa* | 1.483775 | 2.02E-27 |
| *Cd74* | 1.574517 | 1.16E-22 |
| *Gbp4* | 1.127162 | 1.23E-21 |
| *Gbp2* | 1.442935 | 3.36E-19 |
| *Igtp* | 1.08015 | 2.13E-17 |
| *H2-Ab1* | 1.277568 | 1.37E-16 |
| *Ctss* | 1.372639 | 3.69E-16 |
| *Ubd* | 1.905291 | 9.22E-15 |
| **Down regulated** |  |  |
| *Rpl9* | -0.7283 | 2.97E-16 |
| *Cxcl12* | -0.84167 | 6.72E-16 |
| *Rpl21* | -0.58083 | 7.74E-14 |
| *Rpl13* | -0.62302 | 2.34E-13 |
| *Cst3* | -0.69511 | 1.61E-11 |
| *Rps14* | -0.43341 | 2.62E-09 |
| *Rpl18* | -0.563 | 1.25E-08 |
| *Rps15a* | -0.51256 | 1.51E-08 |
| *Rpl30* | -0.51597 | 1.72E-08 |
| *Rps18* | -0.51715 | 2.05E-08 |

**B** **LEC**
| Gene name | Avg. log-fold change | Adj. p value |
| --- | --- | --- |
| **Upregulated** |  |  |
| *B2m* | 1.81363722 | 1.46E-39 |
| *H2-K1* | 1.82143009 | 6.52E-33 |
| *Iigp1* | 2.05863878 | 8.72E-22 |
| *Gbp4* | 1.88908626 | 2.96E-21 |
| *H2-D1* | 1.07551012 | 1.24E-20 |
| *Gbp2* | 1.89596591 | 3.54E-18 |
| *H2-Q7* | 1.26192971 | 3.25E-17 |
| *Gbp7* | 1.31819045 | 3.86E-17 |
| *H2-Aa* | 1.24429821 | 3.81E-15 |
| *H2-Ab1* | 1.1254732 | 1.38E-13 |
| **Down regulated** |  |  |
| *Rpl21* | -0.82606 | 1.30E-25 |
| *Rpl13* | -0.86409 | 6.46E-24 |
| *Rpl30* | -0.74324 | 4.82E-19 |
| *Rps23* | -0.71284 | 2.88E-18 |
| *Rps18* | -0.74298 | 5.32E-18 |
| *Rps11* | -0.62415 | 4.34E-14 |
| *Rpl9* | -0.78012 | 5.04E-14 |
| *Rps25* | -0.68713 | 7.72E-13 |
| *Rps4x* | -0.54557 | 1.19E-12 |
| *Rpl11* | -0.63287 | 3.41E-12 |

In addition, many chemokines were differentially modulated in myeloid cells and ECs with *L. major* infection (Fig 7). Specifically, *Cxcl9* was significantly elevated in macrophages, resident macrophages, inflammatory monocytes, DCs, BECs, and LECs following *L. major* infection (Fig 7 and Fig S3). In contrast, we found significant downregulation of *Ccl24* in resident macrophages, *Cxcl12* in BECs, and *Ccl17* in DCs following *L. major* infection (Fig 7B, 7E and Fig S3). Also important in immune cell recruitment, selectins (*Selp, Sele*) and adhesion molecules (*Vcam1)* were significantly upregulated in BECs with infection, while tight junction molecules like *Cldn5* were downregulated (Fig 7E). Of note, known canonical markers were significantly elevated with *L. major* infection including *Arg1, Nos2,* and *Pla2g7* in macrophages, *Fcgr4, C3,* and *Ccl8* in resident macrophages, and *Ifitm1, Syngr2, Cd200, Ccr7* in DCs (data not shown). The complete list of DEGs that are enriched during *L. major* infection from other cell type clusters such as fibroblasts, keratinocytes, chondrocytes, sebaceous glands, basophils, upper hair follicle cells, pericytes, schwann cells, mast cells, myocytes and myoblasts can be found in Gene Expression Omnibus database with the GEO accession number - GSE181720.

### Characterization of upstream gene regulators and canonical pathways during *L. major* infection in vivo

To determine the cellular and biological mechanisms at the molecular level during *L. major* infection, IPA analysis was performed to define the gene signature for each individual cell type at the site of infection. IPA analysis of our scRNA-Seq data revealed several known and unknown canonical pathways, upstream regulators, and disease-based functional networks. Here, we present the genes that are significantly altered (adj. p value < 0.05) in macrophages, BECs, and LECs from infected ears compared to naive controls.

#### Upstream gene regulators

Our IPA analysis on macrophages, BECs, and LECs revealed potential transcription factors as well as transcriptional targets like anti- and pro-inflammatory genes. In macrophages, we observed 651 upstream regulators in total which include 38 upregulated and 17 downregulated gene regulators. We found cytokines like *IFNγ, IL-4, IL-13, IFNβ1, TNFα*, and transcriptional regulators such as *HIF1α, STAT1, CTCF, TP73, IRF1, MXD1, ATF4, SPI1* mediate macrophage activation upon infection (Table 5). In contrast, transcriptional regulators like *MLXIPL, MYC, TP53, MYCL, CEBPB, GATA1* were inhibited and no cytokines were identified to downregulate macrophages activation following infection (Table 5). We identified 32 regulators were activated and 64 regulators were inhibited with infection in BECs. The upregulated cytokines activating BECs included *IFNγ, TNF, IL-2, IL-4,* while *IL-10* downregulates BEC activation (Table 6). Additionally, we found transcriptional regulators that are either activated (*IRF3, STAT1, IRF7, MXD1*) or inhibited (*MLXIPL, MYC, TRIM24, TP53, SIRT1, HSF1, MYCL, GLIS2, CEBPB, NFE2L2*) in BECs following infection (Table 6). Likewise, 212 upstream regulators were detected of LECs including 17 activated and 23 inhibited regulators. Corresponding to BECs, *IFNγ* increases LEC activation, while *IL-10* downregulates LEC activation upon infection (Table 7). In the LECs, we detected activated transcriptional regulators such as *IRF3, STAT1,* and *IRF7*; however, *MLXIPL, MYC, TRIM24, TP53, SIRT1, MYCL, TARDBP, STAT6, AIRE, LDB1, HNF4A,* and *NFE2L2* were inhibited in LECs comparing infected mice to naive controls (Table 7).

**Table 5.** List of top 5 upstream regulators identified in macrophages by IPA analysis for *L. major* infected ears.

| Predicted activation state | Upstream regulator | Activation z-score | p-value of overlap | No of target molecules in dataset |
| --- | --- | --- | --- | --- |
| **Cytokines** |  |  |  |  |
| Activated | *IFNG* | 4.44 | 9.87E-19 | 36 |
|  | *IL4* | 2.433 | 1.31E-11 | 28 |
|  | *IL13* | 2.713 | 8.71E-09 | 11 |
|  | *IFNβ1* | 2.383 | 0.0000127 | 10 |
|  | *TNF* | 3.364 | 0.0000372 | 12 |
| Inhibited | Not detected | - | - | - |
| **Transcription regulators** |  |  |  |  |
| Activated | *HIF1A* | 2.075 | 1.62E-08 | 13 |
|  | *STAT1* | 2.396 | 6.09E-08 | 12 |
|  | *CTCF* | 2.236 | 5.05E-06 | 5 |
|  | *TP73* | 2.333 | 5.53E-06 | 10 |
|  | *IRF1* | 2.383 | 0.0000841 | 6 |
| Inhibited | *MLXIPL* | -8.062 | 1.2E-88 | 65 |
|  | *MYC* | -7.457 | 3.77E-82 | 71 |
|  | *TP53* | -2.227 | 1.77E-19 | 51 |
|  | *MYCL* | -2.97 | 7.63E-07 | 9 |
|  | *CEBPB* | -2.54 | 0.0000174 | 16 |

**Note:** The activation Z-Scores (-8.062 to 4.44) and *p*-values were < 0.05.

**Table 6.** List of top 5 upstream regulators identified in BECs by IPA analysis for L. major infected ears.

| Predicted activation state | Upstream regulator | Activation z-score | p-value of overlap | No of target molecules in dataset |
| --- | --- | --- | --- | --- |
| **<u>Cytokines</u>** |  |  |  |  |
| Activated | *IFNG* | 3.684 | 5.19E-14 | 23 |
|  | *TNF* | 2.99 | 5.79E-10 | 14 |
|  | *IL-2* | 2.449 | 0.000141 | 7 |
|  | *IL-4* | 2.219 | 0.00156 | 10 |
| Inhibited | *IL-10* | -2.01 | 1.35E-11 | 14 |
| **<u>Transcription regulators</u>** |  |  |  |  |
| Activated | *IRF3* | 3.065 | 2.11E-09 | 10 |
|  | *STAT-1* | 2.377 | 1.37E-08 | 10 |
|  | *IRF7* | 2.573 | 3.74E-07 | 7 |
|  | *MXD1* | 2.0 | 0.00129 | 4 |
| Inhibited | *MLXIPL* | -5.881 | 3.7E-48 | 36 |
|  | *MYC* | -5.207 | 3.54E-43 | 38 |
|  | *TRIM24* | -2.879 | 9.58E-12 | 12 |
|  | *TP53* | -3.877 | 1.8E-11 | 28 |
|  | *SIRT1* | -3.357 | 1.73E-10 | 15 |

**Note:** The activation Z-Scores (-5.881 to 3.684) and p-values were < 0.05.

**Table 7.** List of top 5 upstream regulators identified in LECs by IPA analysis for *L. major* infected ears.

| Predicted activation state | Upstream regulator | Activation z-score | p-value of overlap | No of target molecules in dataset |
| --- | --- | --- | --- | --- |
| **<u>Cytokines</u>** |  |  |  |  |
| Activated | *IFNG* | 3.644 | 1.62E-07 | 14 |
| Inhibited | *IL-10* | -2.219 | 4.28E-07 | 9 |
| **<u>Transcription regulators</u>** |  |  |  |  |
| Activated | *STAT-1* | 2.766 | 3.47E-12 | 12 |
|  | *IRF3* | 3.266 | 9.08E-12 | 11 |
|  | *IRF7* | 2.795 | 2.97E-09 | 8 |
| Inhibited | *MLXIPL* | -6.97 | 2.47E-80 | 49 |
|  | *MYC* | -6.714 | 1.82E-68 | 49 |
|  | *SIRT1* | -4.338 | 8.72E-17 | 19 |
|  | *TRIM24* | -3.564 | 2.14E-14 | 13 |
|  | *MYCL* | -2.97 | 4.91E-10 | 9 |

**Note:** The activation Z-Scores (-6.97 to 3.644) and p-values were < 0.05.

#### Canonical pathways

IPA analysis highlighted an unknown role for eukaryotic translation initiation factor 2 (EIF2) signaling which includes large ribosomal proteins (Rpl) and small ribosomal proteins (Rps) that were significantly downregulated with infection compared to naive animals (Fig 8). Remarkably, the EIF2 pathway was the top downregulated pathway amongst multiple cell types including macrophages (Ranked as 1 amongst 377), BECs (Ranked as 1 amongst 257), and LECs (Ranked as 1 amongst 145) (Fig 8A, C & E). This was followed by an involvement of mTOR signalling and eIF4/p70S6K signalling in macrophages, BECs, and LECs from infected ears. Alongside the IPA pathway results, the expression of individual transcripts for each of the corresponding pathways is provided for macrophages (Fig 8B), BECs (Fig 8D), and LECs (Fig 8F). These data show the top 10 transcripts in the EIF2 signalling pathway in macrophages, BECs, and LECs, which includes many subunits of ribosomal proteins and these transcripts are mostly downregulated in the infected ears compared to naive controls (Fig 8B, D & F). In contrast, the IPA analysis revealed the antigen presentation pathway was increased with infection, and this pathway was also a common feature of infection being the top elevated pathway for macrophages, BECs, and LECs (Fig 8B, D & F). The antigen presentation pathway was enriched in transcripts such as *B2m, Cd74, H2-K1, H2-Aa, H2-Ab1, H2-Eb1* that were elevated with *L. major* infection in macrophages, BECs, and LECs. In addition to macrophages, BECs, and LECs, we also noted that an involvement of mTOR signalling pathway is consistent among the other immune cell types apart of this study such as inflammatory monocytes, DCs, and CD4^+^ T cells, as mTOR signalling is top 5 in the list of pathways (Fig S5). In summary, EIF2 signaling is the top downregulated pathway in BECs, LECs, macrophages, as well as other immune cell types from infected ears.

**Figure 8:**
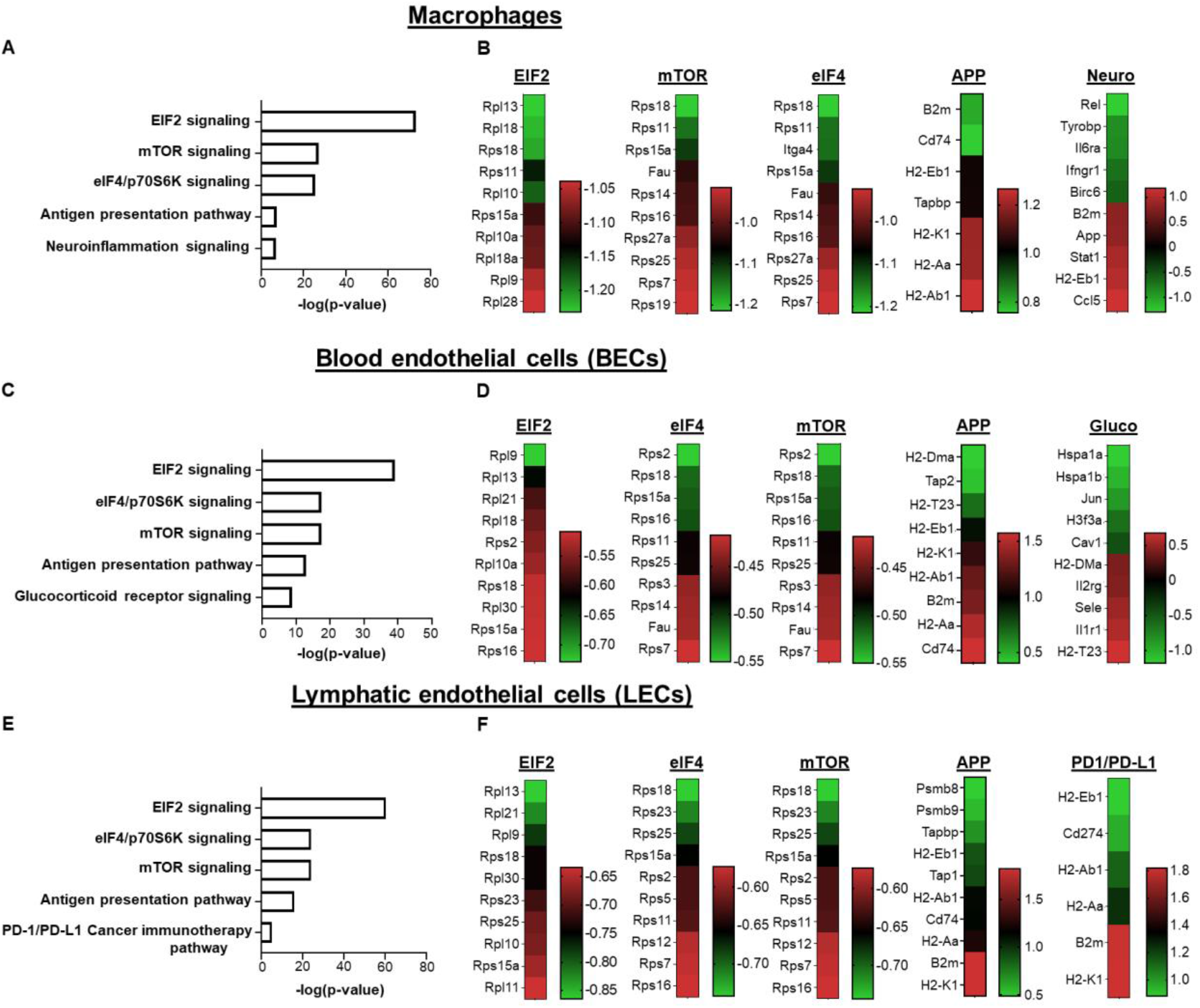
Signaling pathways and molecular networks within individual cell types predicted during *L. major* infection by Ingenuity Pathway Analysis. (A-B) Top 5 differentially regulated canonical pathways and their individual heat maps in macrophages following *L. major* infection. **(C-D)** Top 5 differentially regulated canonical pathways and their individual heat maps in BECs. **(E-F)** Top 5 differentially regulated canonical pathways and their individual heat maps in LECs. The color intensity represents the degree of expression. A red-green color scale was used to reflect the standardized gene expression with red representing high expression and green representing low expression. Cut-off values are adjusted p-value < 0.05.

## Discussion

We conducted a comprehensive high-resolution transcriptomic analysis using both bulk and scRNA-Seq approaches to discover the global changes in the gene expression that occurs following *L. major* infection in vivo. Through our bulk RNA-Seq analyses, we identified many differentially regulated novel transcripts in immune compartments that are related to host immune response pathways. We found significant enrichment of DEGs in the antigen processing and presentation pathway following *L. major* infection. Specifically, our data indicate that *L. major* infection upregulates many MHC molecules belonging to the antigen processing and presentation pathway along with inflammatory cytokines such as *IFNγ* following *L. major* infection. These findings are consistent with the well-established role of the Th1 immune response in parasite control; we confirm antigen presentation is a critical process in host defense to *Leishmania* infection(48–53). Additionally, we noted the enrichment of DEGs specific for other host immune response pathways such as chemokine signalling, cell adhesion molecules, and cytokine-cytokine receptor interactions in infected ears. In contrast, our bulk RNA-Seq results revealed DEGs associated with the ribosomal pathway were downregulated following *L. major* infection. While the importance of the downregulation of transcripts that encode 40S and 60S ribosomal subunits has not been studied in *Leishmania*, these findings suggest cells at the site of infection are actively controlling translation and/or ribogenesis or undergoing a stress response similar to bacterial, viral infection and cancer(54–56). Altogether, our bulk RNA-Seq results confirm a known role for the importance of antigen presentation and highlight an unknown feature of downregulating ribosomal subunits in CL.

Our scRNA-Seq analysis revealed many novel transcripts from various cell types that remain largely unexplored at the site of *L. major* infection. Through our scRNA-Seq data, we defined 35 distinct cell type populations by using canonical markers specific for different cell types including both resident and recruited cells following *L. major* infection. In agreement with previously published results(10, 20), our scRNA-Seq analysis confirmed a significant increase in various immune cell types at the site of infection with recruited myeloid cells such as neutrophils, inflammatory monocytes, and macrophages in lesions (Fig. 6). The transcriptional signature of interferon-induced GTPases like GBPs were significantly upregulated in macrophages, resident macrophages, inflammatory monocytes, DCs, BECs, and LECs following *L. major* infection. These data suggest GBPs may play a protective role in both immune and nonimmune cells during *L. major* infection. GBPs are involved in controlling intracellular pathogen replication and specifically mediating the protection against intracellular pathogens such as *Listeria monocytogenes* and *Mycobacterium bovis*(57) (58). Importantly, mice deficient in GBP genes are more susceptible to *Toxoplasma gondii*(59). Moreover, GBPs restrict *L. donovani* growth in nonphagocytic cells such as murine embryonic fibroblasts in an IFNγ-dependent manner(60).

We detected several transcripts for chemokines (*Cxcl9, Cxcl10*) in myeloid cells that are elevated with infection that may mediate myeloid cell accumulation at the site of infection. Our results revealed a significant downregulation of *Ccl24* transcript in resident macrophages from infected ears compared to naive controls. A recent study demonstrated dermal tissue-resident macrophages shift toward a pro-inflammatory state in *L. major*-infected mice lacking IL-4/IL-13 from eosinophils, which is mainly regulated by CCL24 from resident macrophages(61). Also, potentially mediating myeloid cell migration, we found BECs in infected skin express elevated transcripts for selectins (*Sele, Selp*) and adhesion molecules (*Vcam1*), while concomitantly downregulating transcripts responsible for junctional stability. Mice doubly deficient in E- and P-selectin develop significantly less inflammation following *L. major* infection(62). Taken with our findings, these data suggest BECs play an active role during CL to recruit immune cells into the site of infection. Indeed, these scRNA-Seq data also correlate with our bulk RNA-Seq results which indicate a role of leukocyte trans-endothelial migration pathway in infected ears when compared to naive controls. Overall, our scRNA-Seq data reveals cells in leishmanial lesions exist in pro- inflammatory environment, but the actual host protective role of individual DEGs within each cell type requires further investigation.

We detected the presence of *L. major* transcripts in multiple cell types including myeloid cells like macrophages, neutrophils, and DCs which are all known to harbor parasites(47,63,64). We also detected *L. major* transcripts in stromal cells such as fibroblasts and keratinocytes, which can be infected by parasites(65, 66). Surprisingly, we also found evidence of parasite transcripts in myoblasts, chondrocytes, and BECs which has not previously been reported. Importantly, the molecular techniques used in these studies cannot differentiate between living and dead parasites and we can only report cells harboring parasite transcripts. Therefore, subsequent studies will need to determine if parasites in other non-myeloid cells like myoblasts and BECs are viable and capable of replication and infection. Others have hypothesized *Leishmania* parasites might evade the host immune responses by seeking shelter in different non- macrophage cell types including fibroblasts and keratinocytes in addition to infection in neutrophils, macrophages and DCs which exhibit a more robust pro-inflammatory response against *Leishmania* parasites(67). However, the presence of *L. major* transcripts from non-myeloid stromal cell types like myoblasts and BECs needs to be further explored to determine whether these cells can serve as safe havens for *L. major* parasites during chronic infection or provide a conduit for metastasis. Regardless, these results shed light on cell types previously not thought to harbor *Leishmania* parasites in the skin and provide an opportunity for new investigation.

Macrophages are the replicative niche for the parasite as well as the major cell type responsible for parasite control. BECs play a crucial role in regulating immune cell entry into inflamed tissues and LECs participate in immune cell migration out of the lesions. As a result of the critical roles of these cell types during *L. major* infection, we focused on identifying the upstream regulators and canonical pathways for macrophages, BECs, and LECs. Surprisingly, the antigen processing and presentation pathway and EIF2 signaling were the most significant pathways in all three cell types (macrophages, BECs, and LECs) within infected ears. The antigen processing and presentation showed a positive activation z score indicating overall upregulation of the pathway, which is consistent with findings in human CL lesions(68). Immunoproteasomes play a critical role in the immune response by degrading intracellular proteins to generate MHCI epitopes for effective antigen presentation(69). While the increased expression of immunoproteasome genes in human lesions caused by *L. braziliensis* infection has been reported(68), our results reveal for the first time that LECs express higher levels of transcripts for immunoproteasomes (*psmb8* and *psmb9*), as well as transcripts involved in the antigen presentation pathway (*Tap1* and *Tapbp*) following *L. major* infection. These data suggest that ECs, and specifically LECs, may play an unknown role in antigen presentation during *L. major* infection, similar to viral infection and vaccination(69, 70).

The EIF2 signaling pathway had the highest negative activation z score of all the pathways indicating overall deregulation during CL. To our knowledge, this is the first study highlighting EIF2 signaling as a novel candidate pathway for leishmaniasis. Eukaryotic initiation factor-2 (EIF2) is a GTP-binding protein, which initiates protein translation by delivering charged initiator met-tRNA onto the ribosome(71). Upon subject to infection-induced cellular stress, EIF2 plays a significant role in attenuating translation initiation by phosphorylation of the alpha subunit of eIF2 leading to immediate shut-off of translation and activation of stress response genes(71). Phosphorylation of eIF2α plays as a rate limiting step as it reduces active eIF2-GTP levels at translation initiation which ultimately results in a global reduction of protein synthesis(71, 72). Our data indicate that impaired EIF2 signaling is linked to the downregulation of many ribosomal subunits in macrophages, BECs, and LEC in *L. major*-infected ears (Fig. 8). The known phenomenon of “protein shut off” has been well described at the molecular level for some viruses(73), but there is little to no evidence documenting this phenomenon during *L. major* infection. Generally, the enhanced host response to viral and bacterial infections depends on the upregulation of EIF2-mediated translational control, thereby reducing general protein synthesis(55,56,73). However, we found EIF2 signaling was downregulated with *L. major* infection. Previously it was demonstrated that *L. major* promotes its survival by downregulating macrophage protein synthesis, which is mainly mediated by host translation repressor 4E-BP1(74). Given the robust production of cytokines and chemokines in leishmanial lesions, global protein synthesis does not seem impacted by *L. major* infection in vivo, but careful analysis of EIF2 signaling has not been performed to date.

Alongside EIF2 signalling, the involvement of mTOR and eIF4/p70S6K signalling in macrophages, BECs, and LECs in infected mice hints at an important role for hypoxia-induced oxidative stress at the site of *L. major* infected ears. Our results indicate the activation of metabolic gene targets like hypoxia-inducible factor (HIF-1α) in macrophages, which is consistent with our previous results and work by the Jantsch lab showing leishmanial lesions are hypoxic(75–77). Therefore, we speculate EIF2 signaling is downregulated as part of the stress response, potentially from hypoxia, but mechanism by which EIF2 signaling is impaired in multiple cell types and how that contributes to pathogen control or the pathogenesis of disease in CL is unknown. Collectively, our transcriptome analysis not only provides the first comprehensive list of gene expression at single-cell resolution, but also highlights a previously unknown role of the highly conserved EIF2 signaling pathway in leishmaniasis. Future analysis by us and others utilizing these datasets will expand our knowledge on the complex immune networks and pathways participating in the host response to *Leishmania* infection.

## Materials and methods

### Animals

Female C57BL/6NCr mice were purchased from the National Cancer Institute. Mice were housed in the Division of Laboratory Animal Medicine at University of Arkansas for Medical Sciences (UAMS) under pathogen-free conditions and used for experiments between 6 and 8 weeks of age. All procedures were performed in accordance with the guidelines of the UAMS Institutional Animal Care and Use Committee (IACUC).

### Parasite Infection in vivo

*Leishmania major* (WHO/MHOM/IL/80/Friedlin) strain was used. Parasites were maintained in vitro in Schneider’s Drosophila medium (Gibco) supplemented with 20% heat-inactivated FBS (Invitrogen), 2 mM L-glutamine (Sigma), 100 U/mL penicillin, and 100 mg/mL streptomycin (Sigma). Metacyclic stationary phase promastigotes were isolated from 4–5 day cultures by Ficoll density gradient separation (Sigma). For ear dermal infections, 2×10^6^ *L. major* promastigote parasites in 10 µL PBS (Gibco) were injected intradermally into the ear. Lesion development was monitored weekly by measuring ear thickness and lesion area with a caliper. Lesion volume was calculated. At 4-week post infection, ears were excised and enzymatically digested using 0.25 mg/mL Liberase (Roche) and 10 mg/mL DNase I (Sigma) in incomplete RPMI 1640 (Gibco) for 90 min at 37°C. After digesting, ears were minced manually to obtain cellular content in single cell suspensions. Parasite burdens were determined by limiting dilution assays (LDAs), as previously described(78).

The total RNA was isolated from the cell lysate of naive and infected ears by using Qiagen’s RNeasy plus mini kit according to the manufacturer’s instructions. The CTPR Genomics and Bioinformatics Core at the Arkansas Children’s Research Institute (ACRI) prepared sequencing libraries from RNA samples by use of the Illumina TruSeq Stranded mRNA Sample Preparation Kit v2. for sequencing on the NextSeq 500 platform using Illumina reagents. The quality and quantity of input RNA was determined using the Advanced Analytical Fragment Analyzer (AATI) and Qubit (Life Technologies) instruments, respectively. All samples with RQN (RNA quality number) values of 8.0 or above were processed for sequencing. Sequencing libraries were prepared by use of the TruSeq Stranded mRNA Sample Prep Kit (Illumina). Briefly, total RNA (500 ng) was subjected to polyA selection, then chemically fragmented and converted to single-stranded cDNA using random hexamer primed reverse transcription. The second strand was generated to create double-stranded cDNA, followed by fragment end repair and addition of a single A base on each end of the cDNA. Adapters were ligated to each fragment end to enable attachment to the sequencing flow cell. The adapters also contain unique index sequences that allow the libraries from different samples to be pooled and individually identified during downstream analysis. Library DNA was PCR amplified and enriched for fragments containing adapters at each end to create the final cDNA sequencing library. Libraries were validated on the Fragment Analyzer for fragment size and quantified by use of a Qubit fluorometer. Equal amounts of each library was pooled for sequencing on the NextSeq 500 platform using a high output flow cell to generate approximately 25 million 75 base reads per sample.

### Bulk RNA-Seq: Data Analysis

Following demultiplexing, RNA reads were checked for sequencing quality using FastQC (http://www.bioinformatics.babraham.ac.uk/projects/fastqc) and MultiQC(79) (version 1.6). The raw reads were then processed according to Lexogen’s QuantSeq data analysis pipeline with slight modification. Briefly, residual 3’ adapters, polyA read through sequences, and low quality (Q < 20) bases were trimmed using BBTools BBDuk (version 38.52) (https://sourceforge.net/projects/bbmap/). The first 12 bases were also removed per the manufacture’s recommendation. The cleaned reads (≥ 20bp) were then mapped to the mouse reference genome (GRCm38/mm10/ensemble release-84.38/ <u>GCA_000001635.6</u>) using STAR(80) (version 2.6.1a), allowing up to 2 mismatches depending on the alignment length (e.g. 20-29bp, 0 mismatches; 30- 50bp, 1 mismatch; 50-60+bp, 2 mismatches). Reads mapping to > 20 locations were discarded. Gene level counts were quantified using HTSeq (htseq-counts)(81) (version 0.9.1) (mode: intersection-nonempty).

Genes with unique Entrez IDs and a minimum of ∼2 counts-per-million (CPM) in 4 or more samples were selected for statistical testing. This was followed by scaling normalization using the trimmed mean of M-values (TMM) method(82) to correct for compositional differences between sample libraries. Differential expression between naive and infected ears was evaluated using limma voomWithQualityWeights(83) with empirical bayes smoothing. Genes with Benjamini & Hochberg(84) adjusted p-values ≤ 0.05 and absolute fold-changes ≥ 1.5 were considered significant.

Gene Set Enrichment Analysis (GSEA) was carried out using Kyoto Encyclopedia of Genes and Genomes (KEGG) pathway databases and for each KEGG pathway, a p-value was calculated using hypergeometric test. Cut-off of both *p* < 0.05 and adjusted p-value/FDR value < 0.05 was applied to identify enriched KEGG pathways. DEGs that are more than 1.5-fold in *L. major*-infected ears relative to uninfected controls were used as input, with upregulated and downregulated genes considered separately. Subsequently, the heat maps were generated using these genes with complex Heatmap. All analyses and visualizations were carried out using the statistical computing environment R version 3.6.3, RStudio version 1.2.5042, and Bioconductor version 3.11. The raw data from our bulk RNA-Seq analysis were deposited in Gene Expression Omnibus (GEO accession number - GSE185253).

### scRNA-Seq Sample Preparation

The Genomics and Bioinformatics Core at the Arkansas Children’s Research Institute (ACRI) prepared NGS libraries from fresh single-cell suspensions using the 10X Genomics NextGEM 3’ assay for sequencing on the NextSeq 500 platform using Illumina SBS reagents. The quantity and viability of cells input to the assay was determined using Trypan Blue exclusion under 10X magnification. Library quality was assessed with the Advanced Analytical Fragment Analyzer (Agilent) and Qubit (Life Technologies) instruments.

### scRNA-Seq Data Analysis

Demultiplexed fastq files generated by the UAMS Genomics Core were analyzed with the 10x Genomics Cell Ranger alignment and gene counting software, a self-contained scRNA-Seq pipeline developed by 10X Genomics. The reads are aligned to the mm10 and *Leishmania major* reference transcriptomes using STAR and transcript counts are generated(80, 85) .

The raw counts generated by *cellranger count* were further processed using the R package *Seurat*(86, 87). Low quality cells, potential cell doublets, and cells with high percentage of mitochondrial genes were filtered out of the data. We filtered cells that have unique feature counts over more the 75^th^ percentile plus 1.5 times the interquartile range (IQR) or less than the 25^th^ percentile minus 1.5 time the IQR. Additionally, we filtered cells with mitochondrial counts falling outside the same range with respect to mitochondrial gene percentage. Following filtering the counts all 8 sequencing runs were merged. The counts are then normalized using the LogNormalize method, which normalizes the feature expression measurements for each cell by the total expression, multiplies this by a scale factor (10,000 by default), and log-transforms the result. Next, the 2000 highest variable features are selected. The data is then scaled, and linear regression is included to remove variation associated with percent mitochondria and cell cycle status. Principle component analysis (PCA) is performed on the scaled data. A JackStraw procedure was implemented to determine the significant PCA components that have a strong enrichment of low p-value features.

A graph-based clustering approach is applied(88). Briefly, these methods embed cells in a graph structure - for example a K-nearest neighbor (KNN) graph, with edges drawn between cells with similar feature expression patterns, and then attempt to partition this graph into highly interconnected ‘quasi-cliques’ or ‘communities’. t-distributed stochastic neighbor embedding (tSNE) and Uniform Manifold Approximation and Projection (UMAP)(89) are non-linear dimensional reduction techniques used to visualize and explore the results and are performed using Seurat. Seurat *FindNeighbors* and *FindClusters* functions will be optimized to label clusters based on the visual clustering in the projections. Seurat *FindAllMarkers* function finds markers that define clusters by differential expression. It identifies positive markers of a single cluster compared to all other cells and outputs the differential expression results. These markers will be compared to known markers of expected cell types and results from previous single-cell transcriptome studies in order to assign appropriate cell type labels. Cell type determinations were made by manually expecting these results and some clusters were combined if their expression was found to be similar. Differential expression analysis will be performed using MAST, a GLM-framework that treats cellular detection rate as a covariate(90). The raw data from our scRNA-Seq analysis were deposited in Gene Expression Omnibus (GEO accession number - GSE181720)

### Ingenuity Pathway Analysis (IPA)

QIAGEN’s Ingenuity Pathway Analysis (IPA, QIAGEN Redwood City, www.qiagen.com/ingenuity) was utilized to investigate the DEGs at the level of biochemical pathways and molecular functions. We submitted our DEGs to functional analysis with IPA, and IPA provided canonical pathways, diseases and function, and upstream regulators based on the experimentally observed cause-effect relationships related to transcription, expression, activation, molecular modification, etc. Z-score analyses are used to assess the match between observed and predicted up and down regulation patterns allowing for Bayesian scoring of the results.

### Flow Cytometry Analysis with Statistics

The recruitment of immune cells during *L. major* infection was analyzed by flow cytometry. Cells from both naive and infected ears were incubated with fixable Aqua dye (Invitrogen) to assess cell viability, and cells were treated with FcγR blocking reagent (Bio X Cell) prior to staining for the following markers: anti-CD45-AF700 (clone 30-F11), anti-Ly6C-PerCP-Cy5.5 (clone HK1.4), and anti-Ly6G-eFlour 450 (clone 1A8) were purchased from eBiosciences; anti-CD64-BV711 (clone X54-5/7.1), anti-CD11b-BV605 (clone M1/70), anti-CD31-AF488 (390), and anti-podoplanin-PE/Dazzle 594 (clone 8.1.1) were purchased from BioLegend. Cells were acquired using an LSR II Fortessa flow cytometer (BD Biosciences) and analyzed using FlowJo software version 10.2 (Tree Star). All statistical analyses were performed using Prism version 8.0 (GraphPad Software, Inc.). Comparisons between groups were performed using the two-tailed Students unpaired *t*-test.

## Acknowledgements and funding sources

The authors would like to thank Dr. Lu Huang, the Huang lab, Lucy Fry and Conner Webb for their critical reading of the manuscript. We would like to thank the veterinary staff from the Division of Laboratory Animal Medicine (DLAM) and Andrea Harris from Flow cytometry core facility at UAMS for their excellent technical assistance. This work was supported by the Center for Microbial Pathogenesis and Host Inflammatory Responses (funded by National Institutes of Health National Institute of General Medical Sciences Centers of Biomedical Research Excellence Grant P20-GM103625) and the Oak Ridge Associated Universities (ORAU) 2019 Ralph E. Powe Junior Faculty Enhancement Award to Dr. Weinkopff. This work was also supported by the UAMS Translational Research Institute in collaboration with the UAMS Division for Diversity, Equity and Inclusion (DDEI) Mini Grants for Under-Represented Faculty Researchers to Dr. Weinkopff as part of the NIH National Center for Advancing Translational Sciences (NCATS), UL1-TR003107. This study was supported by the Arkansas Children’s Research Institute, the Arkansas Biosciences Institute, and the Center for Translational Pediatric Research funded under the National Institutes of Health National Institute of General Medical Sciences (NIH/NIGMS) grant P20GM121293 and the National Science Foundation Award No. OIA-1946391. The funders had no role in study design, data analysis, decision to publish or preparation of the manuscript.

## Supporting Information

**Fig S1.**
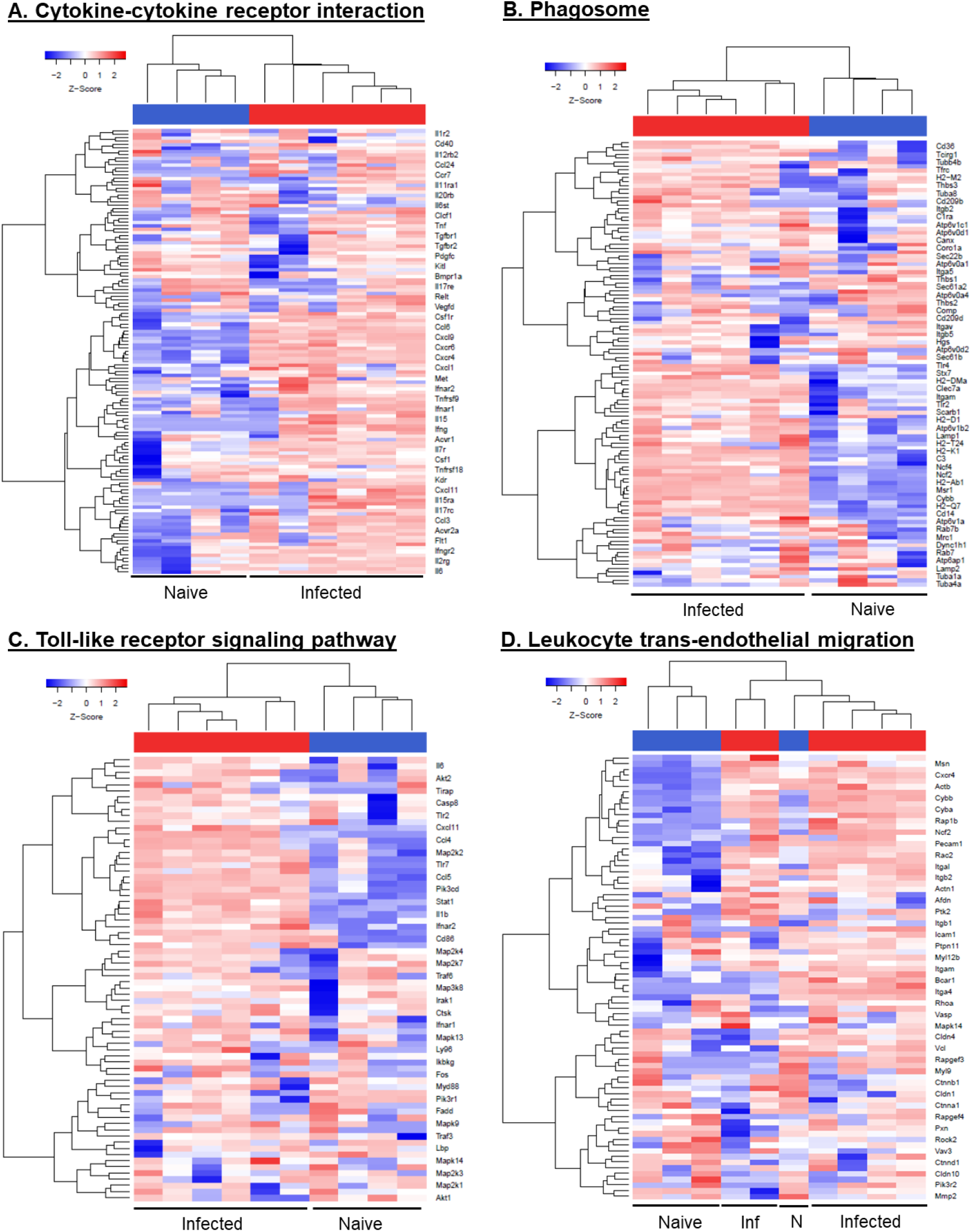
Heat map analysis showing transcriptional responses from other immune-related pathways during *L. major* infection in vivo. The DEGs involved in the other host immune response pathways by KEGG enrichment analysis (A, B, C and D) in the infected ears compared to naïve mice presented as heat maps. Hierarchical clustering of the expression profile was grouped according to functional categories. Heat maps indicate the FC in *L. major* infected ear gene expression >2-fold (red) or <2-fold (blue).

**Fig S2.**
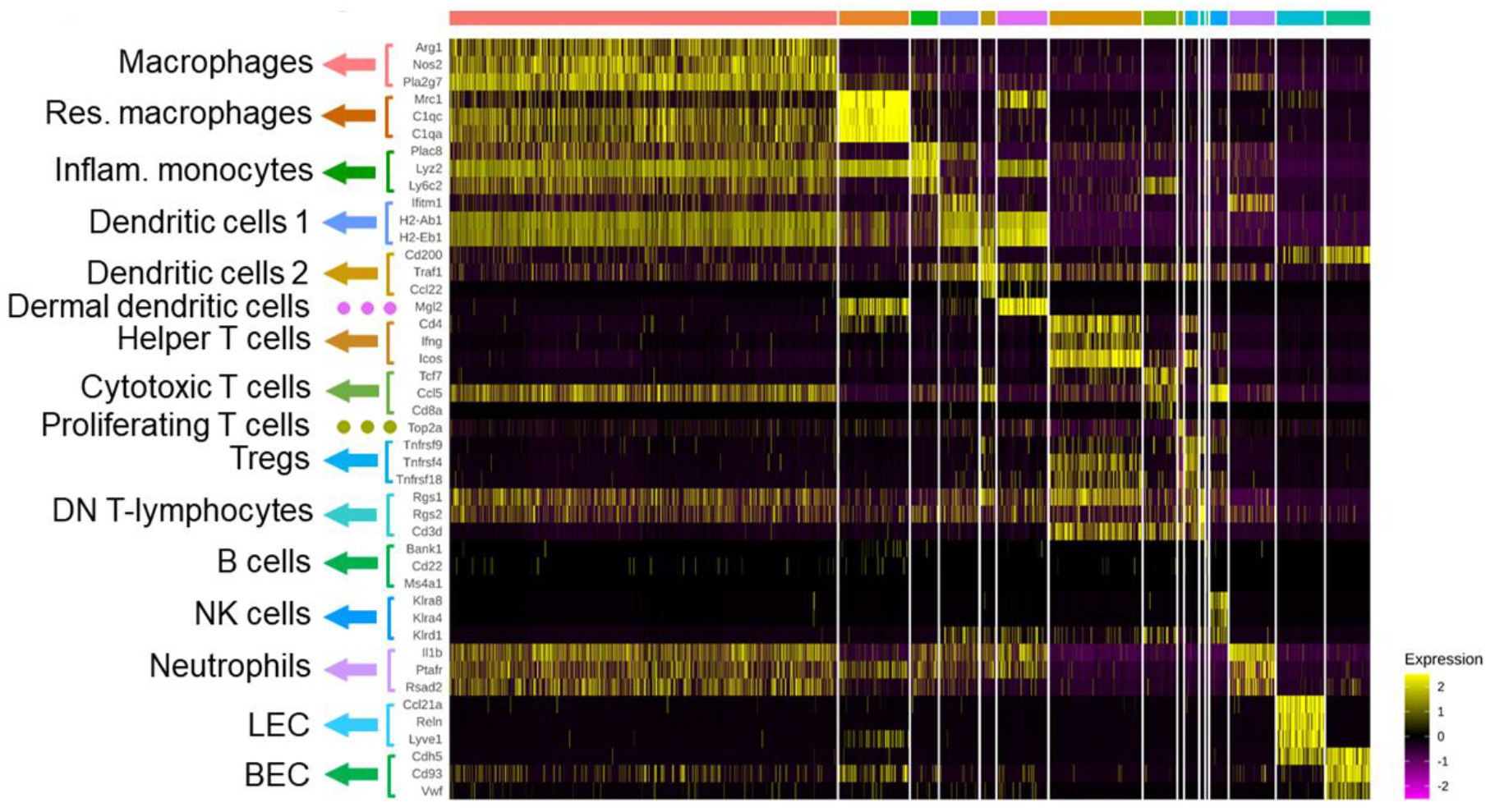
Differentially expressed genes in selected immune cell types during *L. major* infection. Heat map showing the three highly expressed genes for at least 14 immune cell clusters that were selected along with BECs and LECs. Each column represents a single cell and each row represents an individual gene. Three marker genes per cluster was color-coded and shown on the left. Yellow indicates maximum gene expression and purple indicates no expression in scaled log-normalized unique molecular identifier counts.

**Fig S3.**
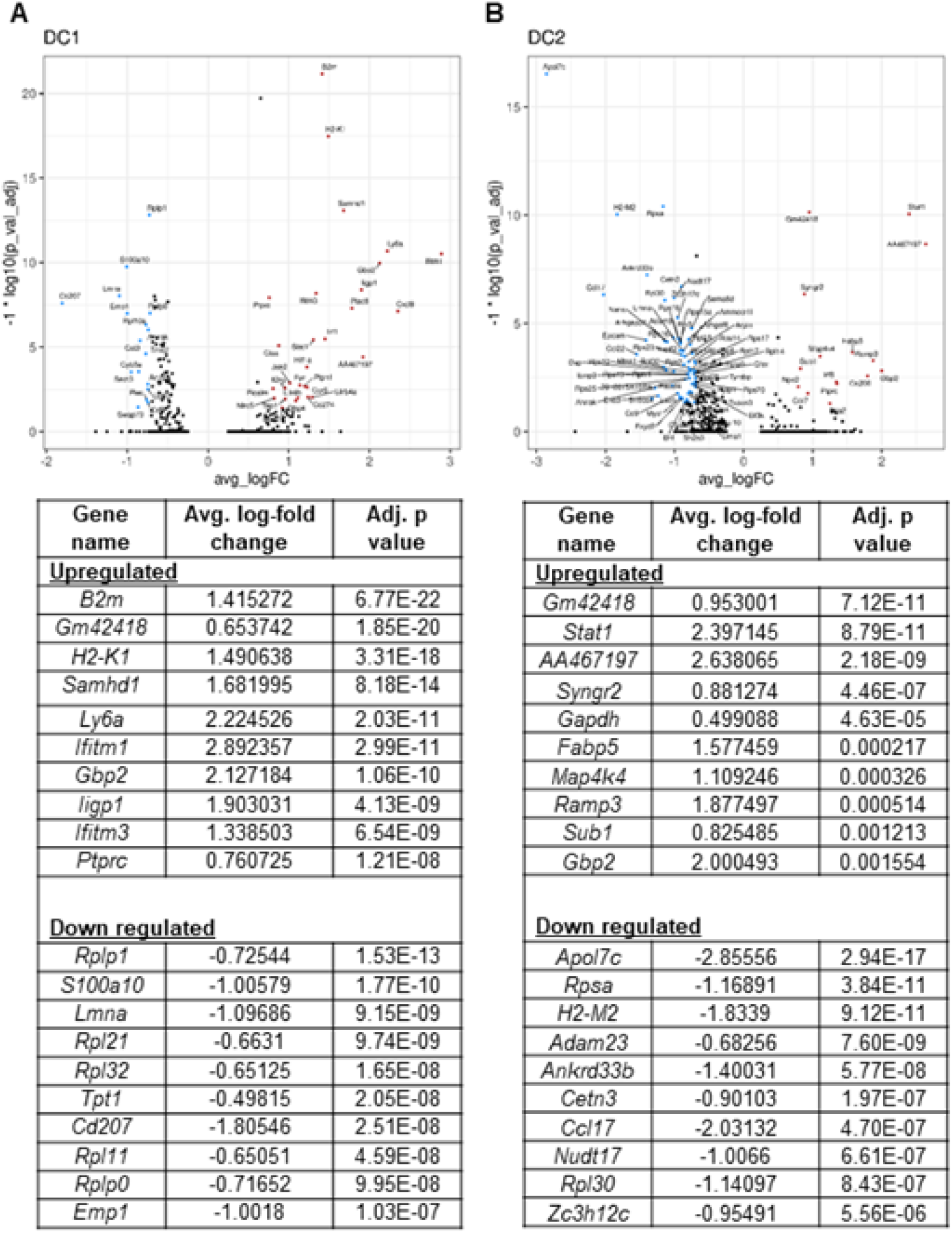
Differentially expressed genes in DCs during *L. major* infection. Volcano plot showing the DEGs in dendritic cells (DC1 and DC2) and list includes the top DEGs enriched in DCs following *L. major* infection. Colored dots indicate genes at least 2 (natural log ∼0.693) fold increased (red) or decreased (blue) in infected cells relative to naïve cells with an adjusted p-value < 0.05.

**Fig S4.**
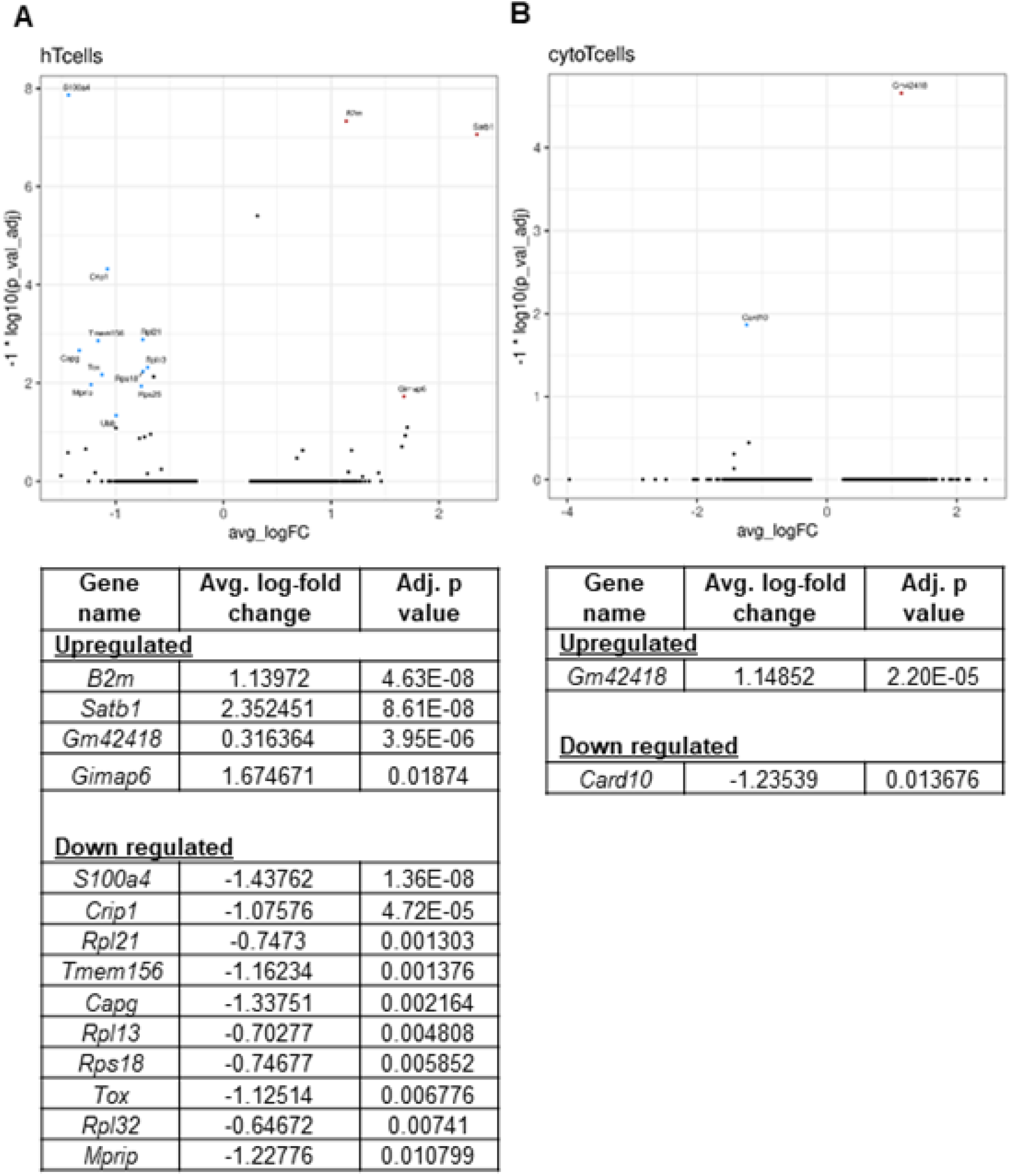
Differentially expressed genes in CD4^+^ and CD8^+^ T cells during *L. major* infection. Volcano plot showing the DEGs in CD4^+^ and CD8^+^ T cells and list includes the top DEGs enriched in CD4^+^ and CD8^+^ T cells following *L. major* infection. Colored dots indicate genes at least 2 (natural log ∼0.693) fold increased (red) or decreased (blue) in infected cells relative to naïve cells with an adjusted p-value < 0.05.

**Fig S5:**
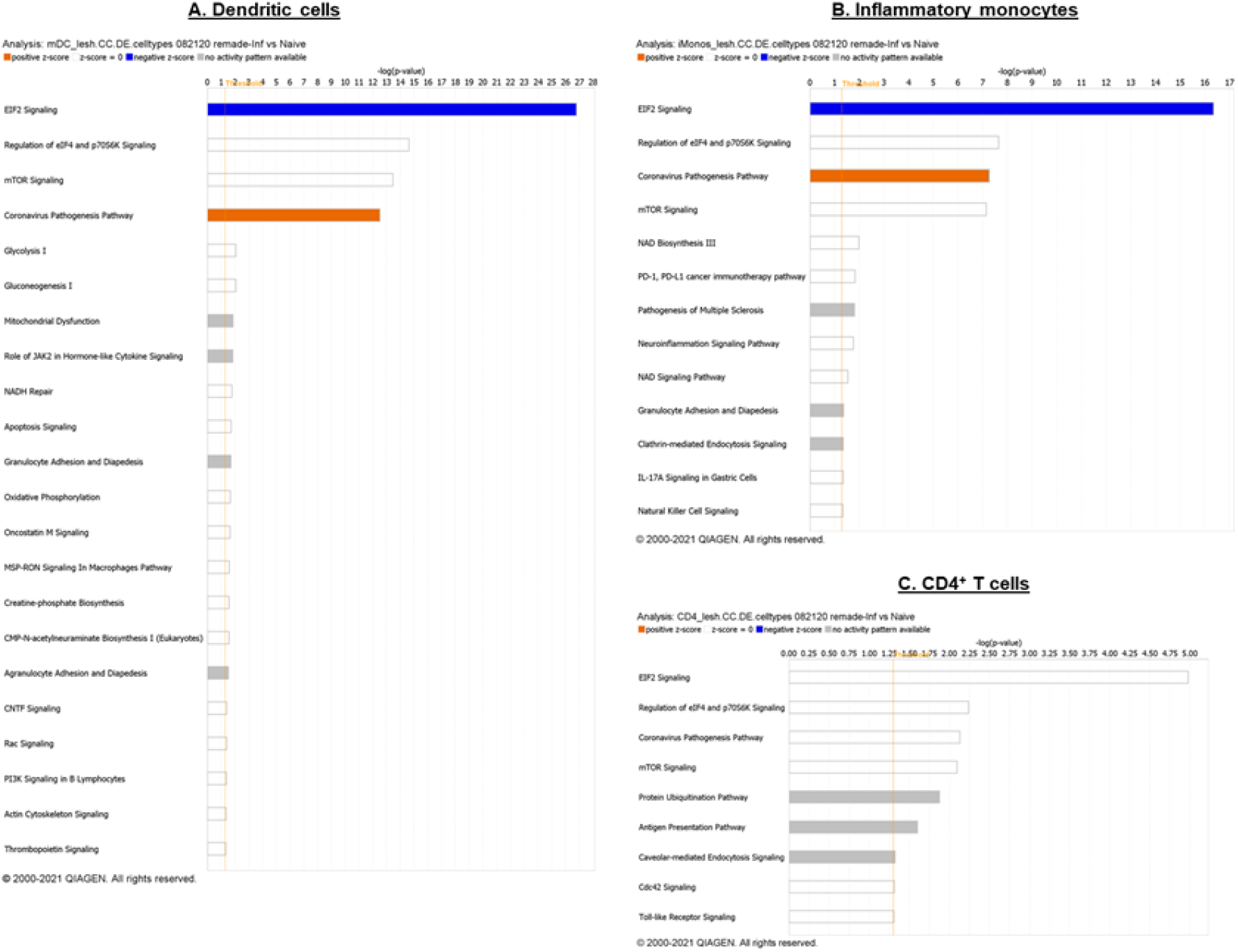
IPA predicted the role of mTOR signaling in other immune cell types during *L. major* infection by. (A-C) Differentially regulated canonical pathways in DCs (A), inflammatory monocytes (B), CD4^+^ T cells following *L. major* infection.

**Table S1.**
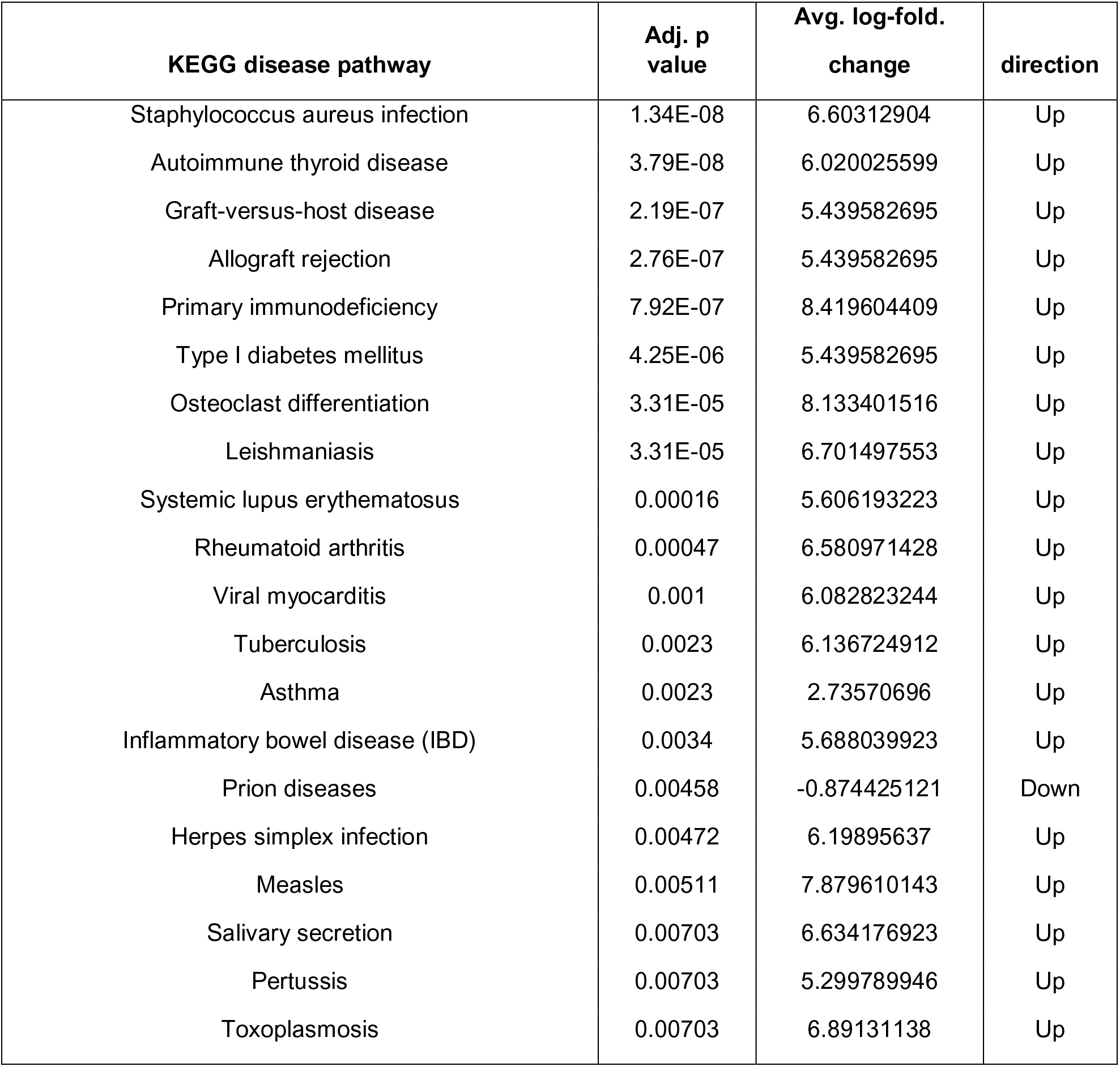
List of top 20 KEGG disease pathways enriched for differentially expressed genes (DEGs).

